# Sacrificing Adaptability for Functionality: The Ivory Tower of Macular Müller Cells

**DOI:** 10.1101/2024.04.28.590478

**Authors:** Ting Zhang, Kaiyu Jin, Shaoxue Zeng, Penghui Yang, Meidong Zhu, Jialing Zhang, Yingying Chen, Sora Lee, Michelle Yam, Yue Zeng, Xiaoyan Lu, Lipin Loo, G. Gregory Neely, Andrew Chang, Fanfan Zhou, Jianhai Du, Xiaohui Fan, Ling Zhu, Mark C. Gillies

**Affiliations:** Macula Research Group, Save Sight Institute, Faculty of Medicine and Health, The University of Sydney, Sydney, NSW 2006, Australia; Pharmaceutical Informatics Institute, College of Pharmaceutical Sciences, Zhejiang University, Hangzhou, Zhejiang 310058, China; New South Wales Tissue Bank, New South Wales Organ and Tissue Donation Service, Sydney, NSW, 2000, Australia; Department of Ophthalmology, West China Hospital, Sichuan University, Chengdu, Sichuan 610041, China; Charles Perkins Centre, Dr. John and Anne Chong Lab for Functional Genomics, Centenary Institute and School of Life and Environmental Sciences, University of Sydney, Camperdown, New South Wales, Australia; Save Sight Institute, Faculty of Medicine and Health, The University of Sydney, Sydney, NSW, 2000, Australia; Molecular Drug Development Group, Sydney Pharmacy School, Faculty of Medicine and Health, The University of Sydney, Sydney, NSW 2006, Australia; Departments of Ophthalmology and Visual Sciences and Biochemistry and Molecular Medicine, West Virginia University, Morgantown, WV, 26506, United States; National Key Laboratory of Chinese Medicine Modernization, Innovation Centre of Yangtze River Delta, Zhejiang University, Jiaxing 314103, China

## Abstract

The predilection of many retinal diseases for the macula suggests it may be less resistant to stress than the peripheral retina. Profiling of single-cell level transcriptional changes found that the peripheral retina exhibited more transcriptional changes than the macula in response to stress. One pronounced change was in a subgroup of Müller cells (MCs) that were dominant in the peripheral retina. Genes more abundantly expressed in peripheral MCs were mainly associated with stress responses and were more influenced by light stress. In contrast, genes highly expressed in MCs that predominated in the macula had roles in cellular function and were less influenced by light stress. Metallothionein 1, A Kinase Anchor Protein 12 and MAF BZIP Transcription Factor F were more abundantly expressed in peripheral MCs than in macular MCs. Knockdown of these genes in primary MCs reduced their viability under stress. Our findings indicate that macular MCs are more directed toward maintaining retinal function rather than mounting a stress response when exposed to stress, which may contribute to the macula’s vulnerability to degenerative diseases.

## Introduction

The neurosensory retina is a remarkable tissue that converts light stimuli into neurochemical signals that are transmitted to the brain for visual perception. It comprises diverse cell types, including neurons, such as photoreceptors, horizontal cells, bipolar cells, amacrine cells and retinal ganglion cells, and Müller cells which are the main glial cell of the retina. The complex interaction of neurons and glial cells maintains retinal structure and homeostasis, ensuring optimal visual function.

The macula, a specialised cone-rich region located at the centre of the human retina, is responsible for high-acuity central vision. This functionally important area is more prone than the peripheral retina to develop many diseases, such as age-related macular degeneration (AMD), diabetic macular edema (DME) and macular telangiectasia type 2 (MacTel) [1, 2] for reasons that remain largely unknown. We hypothesise that the macula is more prone to developing disease than the peripheral retina because it is less able to mount an effective response against stress.

Müller cells, the principal glial cells of the retina that span the entire thickness of the vertebrate neural retina, play a crucial role in maintaining the structural and functional stability of the retina. They are specifically involved in regulating transcellular ion and water transport, maintaining the blood-retinal barrier and modulating neurotransmitter levels [3]. Additionally, they supply trophic factors, metabolites and antioxidants to photoreceptors and neurons, store glycogen and facilitate light transmission to photoreceptors. Müller cells also can regenerate neurons and photoreceptors in response to stress or injury in non-primate species [4]. We have previously reported that Müller cells that predominate in the macula have distinct transcriptomic profiles, as evidenced by bulk RNA sequencing results, from those that are dominant in the peripheral retina [5]. The differential response to stress of these two populations, if there is one, is yet to be clarified.

Light exposure induces age-related and oxidative stress in the retina through various mechanisms, including generating reactive oxygen species (ROS), chronic inflammation, photoreceptor damage and compromised cellular repair mechanisms [6–8]. Light-stressed retinas in laboratory animals have been widely used in the study of retinal degeneration [8, 9]. In this study, we compared the single-cell transcriptomic profiles of human retinal explants from both the macula and peripheral retina exposed to high-intensity or dim light. We hoped to gain insights into the reasons underlying the macula’s particular vulnerability to developing degenerative diseases.

## Results

### Identification of human retinal cell types in the *ex vivo* cultured macula and peripheral retinas

We conducted a transcriptional analysis of 16 retinal explants from 4 *postmortem* human donor eyes **(Supplementary Table 1)**, comprising paired macular and peripheral retinas with and without light stress. The macular and mid-peripheral neural retinas from the left eyes were exposed to intense light (32K lux) for four hours, while the corresponding regions from the right eyes were subjected to dim light (5 lux) as controls (**Supplementary Figure 1**). We then investigated gene expression at the single-cell level (**Figure 1A)**.

**Figure 1.**
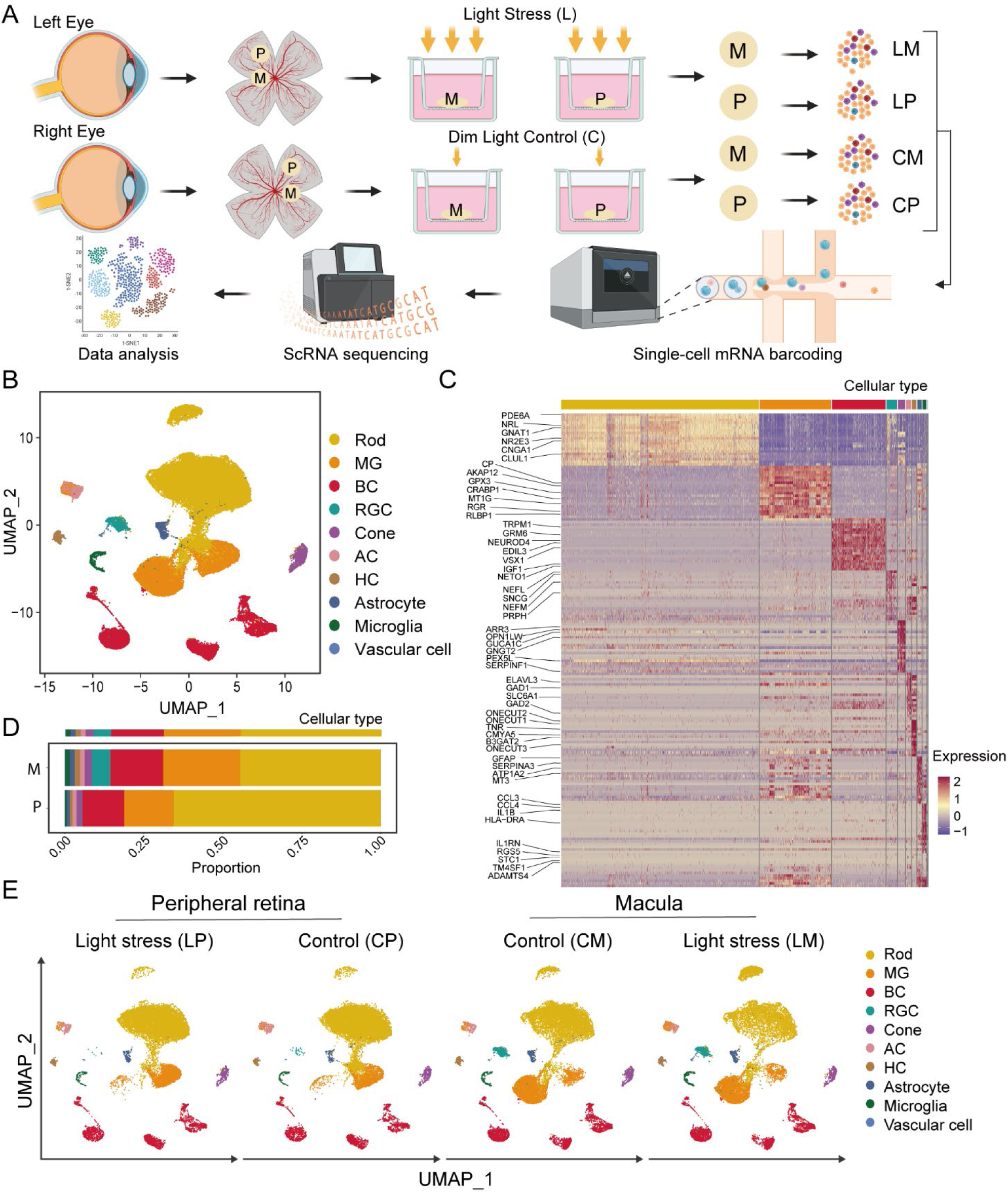
Schematic workflow for generating single-cell retinal cells and transcriptomic profiles of retinal cells dissociated from the macula and mid-peripheral retina by single-cell RNA sequencing analysis. **A.** The workflow illustrates the steps involved in generating single cells from human macular and peripheral retinal explants, including exposure to either bright or dim light for 4 hours. The process begins with the dissection of human donor eyes to isolate the neural retina, followed by culturing the macula and mid-peripheral retinal explants on transwells. The retinal cells are dissociated from the tissue to create a single-cell suspension. n=4 donors per treatment group. **M:** macula, **P:** peripheral retina, **L:** light stress, **C:** dim light, **LM:** macula with light stress; **LP:** peripheral retina with light stress; **CM:** macula with dim light; **CP:** peripheral retina with dim light. **B.** Retinal cell clustering using uniform manifold approximation and projection (UMAP) for all detected transcriptomes. **C.** Heatmap showing the expression of retinal cell markers in identified cell clusters. **D.** Proportions of retinal cell types in the macula and peripheral neural retinas. **E.** UMAP for the peripheral retinas (**P**) and maculas (**M**) without (**C**) or with (**L**) light stress.

We obtained 77,405 transcriptomic profiles, including 19,419 profiles from the <u>P</u>eripheral retinas with <u>L</u>ight stress (**LP**), 19,887 profiles from the <u>P</u>eripheral retinas with dim light as the <u>C</u>ontrol (**CP**), 18,290 profiles from the <u>M</u>acula with <u>L</u>ight stress (**LM**) and 19,809 profiles from the <u>M</u>acula with dim light as the <u>C</u>ontrol (**CM**). The sequencing depth (indicated by the average number of unique genes detected per cell) is illustrated in **Supplementary Figure 2A**. Overall gene expression patterns were very similar across the four donors (**Supplementary Figure 2B**).

We employed unsupervised clustering, uniform manifold approximation and projection (UMAP) dimensionality reduction techniques to analyse the integrated dataset. Based on the expression of established retinal cell-specific markers, we identified ten major retinal cell populations (**Figure 1B** and **Supplementary Figure 2C**). Each cell population was characterised by a set of unique signature genes (**Figure 1C**): rods (42,687 cells), cones (1,573 cells), bipolar cells (11,530 cells), retinal ganglion cells (2,281 cells), amacrine cells (1,071 cells), horizontal cells (965 cells), Müller cells (15,430 cells), astrocytes (943 cells), microglia (814 cells) and vascular endothelial cells (111 cells).

The cellular composition between the macula and peripheral retina was significantly different (**Figure 1D**): retinal ganglion cells (5.87% vs. 0.11%), cones (2.32% vs. 1.75%) and rods (44.5% vs. 65.5%). We also examined the distribution and expression pattern of the cell populations across the four treatment groups (LP, CP, LM and CM) (**Supplementary Figure 2D**). The proportions of cell populations in the macula and peripheral retinas did not differ between the groups exposed to bright or dim light. The degree of transcriptomic changes caused by light stress was not as great as the differences between the macula and peripheral retina (**Figure 1E**).

### Transcriptomic changes between the macular and peripheral retinal cells in response to stress

To examine the differential responses of macular and peripheral retinal cells to light stress, we assessed the gene expression changes across all retinal cells in the four treatment groups (LP, CP, LM and CM). The Venn diagram (**Figure 2A)** illustrates the number of genes significantly altered after light stress in the peripheral retinas (blue circle: 198 genes) and the macula (red circle: 82 genes). 30 genes were differentially expressed in both the peripheral retina and macula in response to light stress, while 742 genes were differentially expressed between the peripheral retina and macula without light stress. The correlation analysis found that the peripheral retina (LP vs. CP) displayed significantly more transcriptomic changes than macula (LM vs. CM) in response to light stress (**Figure 2B**). In this heatmap, a higher correlation (red colour) represents fewer changes.

**Figure 2.**
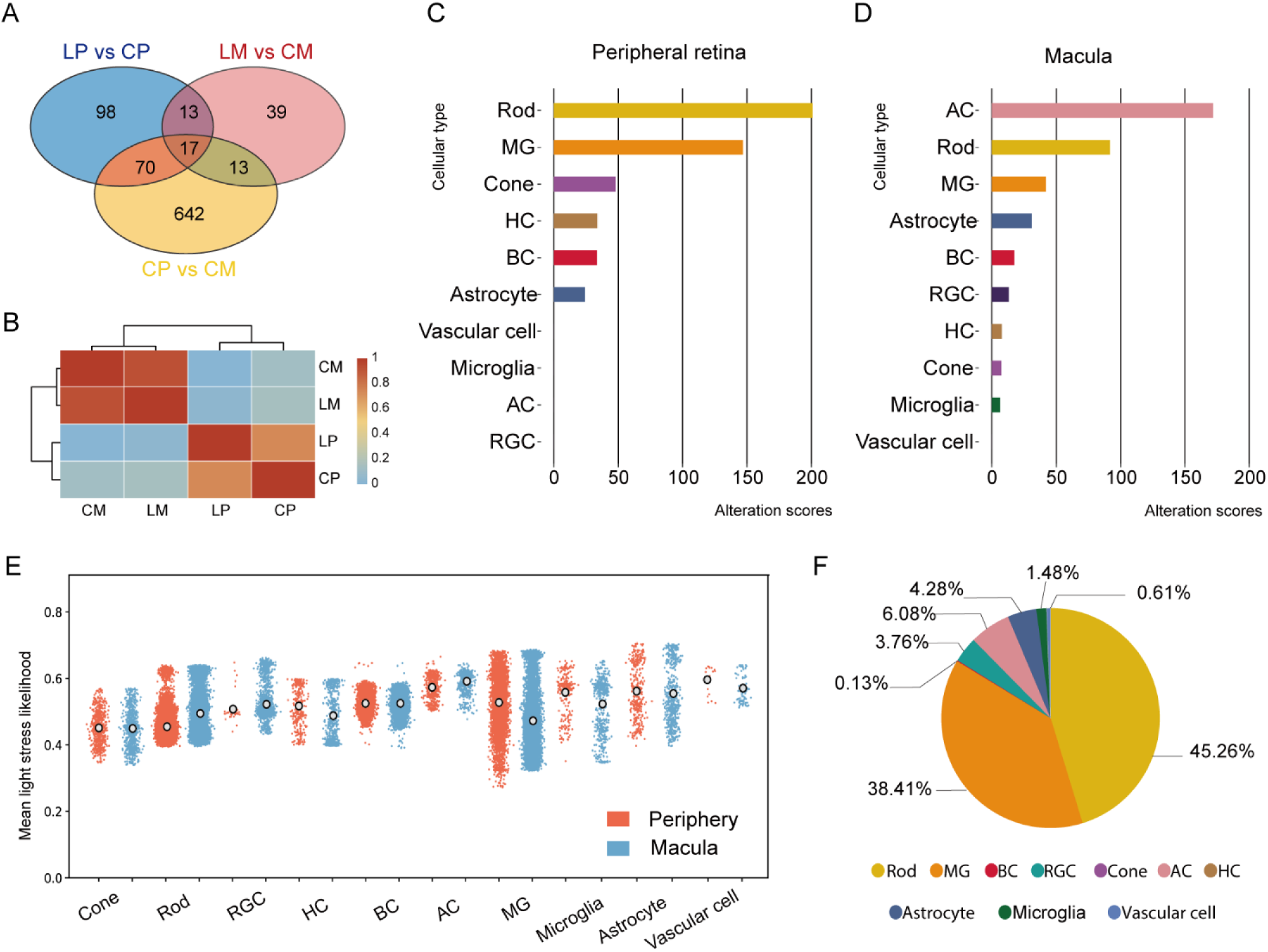
Regional and cell type-specific alterations in response to stress. **A**. A Venn diagram illustrates the relationships among LP vs. CP, LM vs. CM and CP vs. CM. **B**. A heatmap shows the correlations between different treatment groups. High correlation (red) between CM and LM indicates fewer transcriptional changes, while low correlation (orange) between CP and LP suggests significant transcriptomic profile changes. **LP**: Light-stressed Peripheral retina, **CP**: Control Peripheral retina, **LM**: Light-stressed Macula, **CM**: Control Macula. **C** & **D**. Alteration scores for retinal cell types in the peripheral retina (**C**) and macula (**D**) in response to light stress. Higher alteration scores indicate more significant changes. In the peripheral retina, significant changes were observed in rods and Müller glial (MG) cells. In the macula, amacrine cells and rods exhibited the most significant changes. **E**. A scatter plot of all single cells from the macula and peripheral retina showing the distribution of light stress-associated likelihood values. Outlined circles represent average likelihood values. **F**. Proportions of each cell type among all cells from both the macula and periphery with a light stress-associated relative likelihood value greater than 0.6.

We divided the overall changes shown in the heatmap into different clusters of retinal cell types to explore cell-type-specific responses to stress in the peripheral retina (**Figure 2C**) and macula (**Figure 2D)**. We introduced an ‘alteration score’ to measure changes in the quantity and expression levels of signature genes during biological processes, quantifying the extent of variation in these cell types following biological perturbation [10]. Higher alteration scores indicate more significant changes. Peripheral Müller cells and rods had significantly higher alteration scores than macular Müller cells and rods, suggesting that the peripheral retina is more responsive to light-induced stress than the macula. We observed that amacrine cells in the macula were highly sensitive to light stress, with significant and greater transcriptomic changes than amacrine cells in the peripheral retinas. We also observed that hyperglycemic stress induced more transcriptomic changes in the peripheral retina than in the macula, with higher alteration scores in peripheral Müller cells and rods compared to their macular counterparts (**Supplementary Figure 3**). These findings suggest a generalised divergence in stress responses between the macula and peripheral retina.

To further explore the impact of light stress on each cell type, we calculated a light stress-associated relative likelihood (y-axis) for each cell (dots) using a manifold approximation based on all cells across both the dim light control and light stress groups (**Figure 2E**). Examination of the distribution of these values across the ten previously identified cell types revealed that Müller cells had a wider range of light stress-associated relative likelihood values than other cell types (**Figure 2E**). The peripheral Müller cells exhibited a higher mean value of light stress-associated likelihood compared to macular Müller cells, indicating a greater response to light stress. **Figure 2F** displays the proportion of cells from each cell type in all cells from both the macula and periphery that had a high light stress-associated relative likelihood value (> 0.6): these cells were mostly rods or Müller cells.

Given the known sensitivity of rod cells to light stress, understanding their response is pivotal to interpreting the broader retinal reaction. We conducted bioinformatic analyses on rod cells from both the macula and peripheral retina in response to light stress. Seven subpopulations of rod cells were identified (**Supplementary Figure 4A**), with the top three subgroups constituting over 93% of the total rod population. The distribution of these top three subgroups between the macula and peripheral retina was found to be similar (**Supplementary Figure 4B**). We then identified the significantly altered genes in response to light stress within these subpopulations, categorizing them based on their macular or peripheral retinal origin. Ingenuity Pathway Analysis (IPA) revealed that despite differences in the degree of transcriptional changes, the key canonical pathways affected were largely the same, with four out of the top five pathways being identical (**Supplementary Figures 4C-D**).

### Divergent light-induced responses and biological functions of human macular and peripheral Müller cells

We investigated the differentially expressed genes (DEGs) between total Müller cells from the macula and peripheral retina after exposure to light stress and hyperglycemic stress across four treatment groups (**Supplementary Figure 5A-D)**. The volcano plot (**Figure 3A**) depicts the top 40 DEGs that were significantly more highly expressed in peripheral Müller cells compared to macular Müller cells (displayed on the left) and *vice versa* (displayed on the right). We subsequently analysed the modulation of these 80 genes under light-induced stress. Red dots in the volcano plot signify genes that were significantly upregulated (>9%) under light stress, while blue dots indicate those that were significantly downregulated (>9%). We found that 25 of the top 40 highly expressed genes in peripheral Müller cells (left) were significantly differentially expressed in response to light stress (19 genes upregulated and 6 downregulated) in contrast to only 6 of the top 40 highly expressed genes in macular Müller cells (right) (all 6 downregulated).

**Figure 3.**
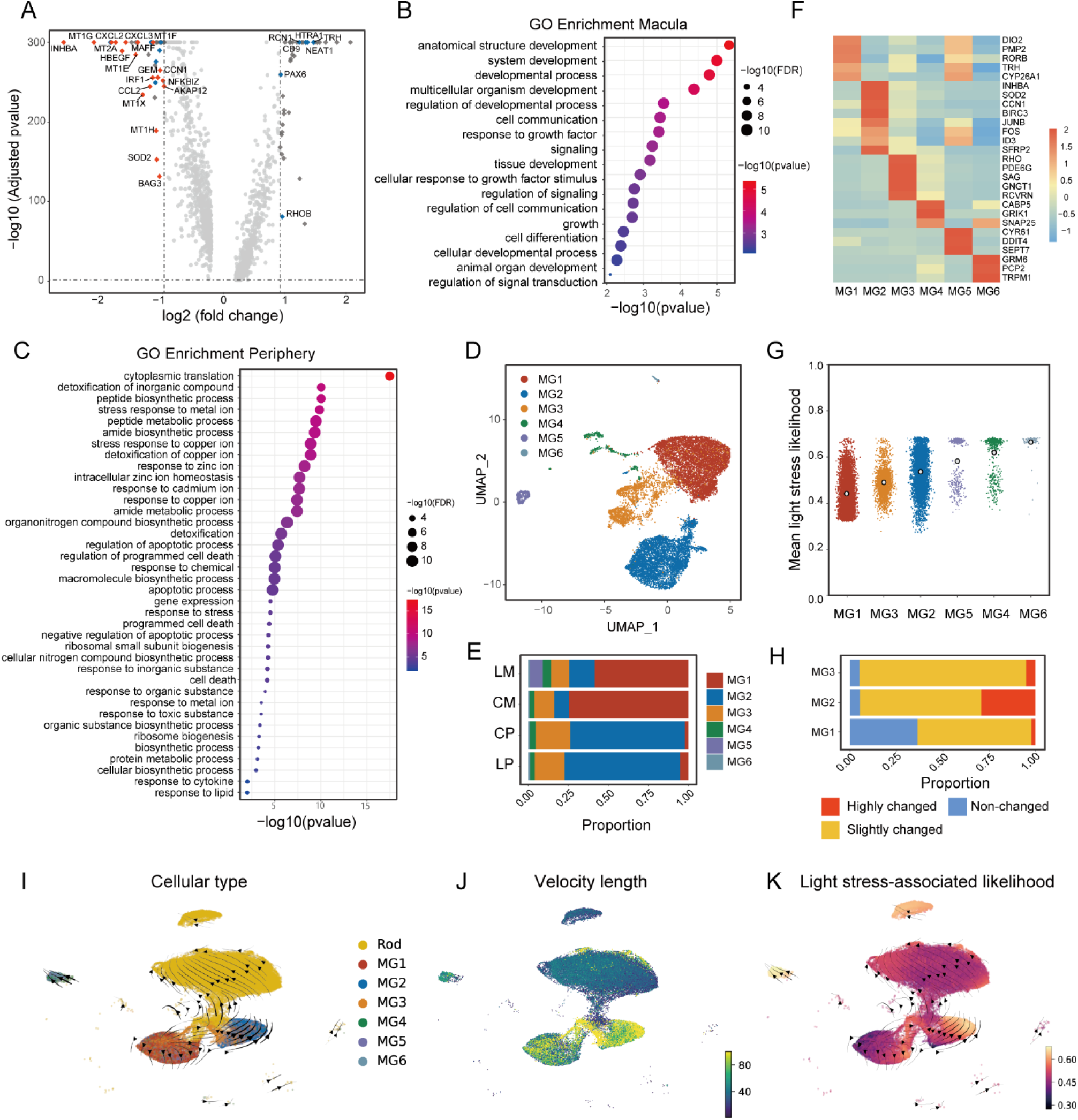
Subclusters of human Müller cells from the macula and peripheral retinas. **A**. Volcano plot of DEGs in Müller cells in the macula and peripheral retinas showing genes that were significantly differentially regulated by light stress. Red dots were upregulated while blue dots were downregulated. **B**. GO analysis of genes highly expressed in macular Müller cells (m-HXs) revealed significant enrichment in biological processes related to cell function and development. **C**. GO analysis of genes highly expressed in peripheral Müller cells (p-HXs) identified significant enrichment in various biological processes related to stress responses. **D**. UMAP of subclusters of human Müller cells identified from the macula and peripheral retinas. **E**. Proportions of Müller cell subclusters in the macula (**M**) and periphery (**P**) with (**L**) and without (C) light stress. **F**. Heatmap of the cluster with marker genes highly expressed in various subtypes of Müller cells. **G**. Scatter plot of six subtypes of Müller cells revealing the likelihood values of transcriptomes upon light stress. Each dot represents one Müller cell. **H**. Proportions of highly-changed (red), slightly-changed (orange) and non-changed (blue) cells in subtypes of Müller cells in response to light stress. **I**. UMAP plot of velocity and trajectory analyses revealed transition dynamics from Müller cells to rods. Colours indicate different cell types. **J**. UMAP plot of velocity length displaying the speed of differentiation. **K**. UMAP plot of single-cell velocity visualized by the previously calculated light stress-associated likelihood values.

We compared the gene expression profiles of macular and peripheral Müller cells to elucidate their functional differences. We identified genes with more than a 1.5-fold expression difference in either macular or peripheral Müller cells, designating them as m-HXs (highly expressed genes in macular Müller cells) and p-HXs (highly expressed genes in peripheral Müller cells), respectively (**Supplementary Data 1**). Subsequent Gene Ontology (GO) analysis of both m-HXs and p-HXs focused on responses to light stress (**supplementary Data 2**). Interestingly, m-HXs did not exhibit significant GO term enrichment related to light stress responses. However, they were significantly enriched in biological processes associated with the development of anatomical structures, cell communication, signaling, differentiation, and signal transduction (**Figure 3B, Supplementary Figure 5E**). Conversely, p-HXs, which were differentially upregulated in peripheral Müller cells in response to light stress, showed enrichment in pathways related to stress responses, including reactions to various compounds, metal ions, stress, cell death, metabolic processes, detoxification, and regulation of cell death (**Figure 3C, Supplementary Figure 5E**). The specific numbers of genes involved in each enriched biological process from the GO analysis are summarized in Supplementary Data 3.

### Different light-induced responses and biological functions of different subtypes of Müller cells

We identified 6 subtypes (MG1 to MG6) of Müller cells in the transcriptomes obtained from the LP, CP, LM and CM groups (**Figure 3D**). The proportion of two major subtypes, macula-dominant Müller glia (MG1) and peripheral retina-dominant Müller glia (MG2), accounted for approximately 75% of Müller cells in the macula and peripheral retinas (**Figure 3E**). We then focused on three subtypes (MG1 to MG3) due to their high abundance; the MG4 subtype contained relatively few cells and the MG5 and MG6 subtypes were donor-specific (**Supplementary Figure 5F**). We generated a heatmap to cluster the top markers that were highly expressed in various Müller cell subtypes (**Figure 3F**). We also cross-referenced the highly expressed feature genes in the different Müller cell subtypes (MG1, 2 and 3) with those reported in previous studies on Müller cells. We found MG1 cells had a high expression of feature genes, such as *COL4A3*, *FGF9*, *DIO2* and *ZNF385D*, in macular Müller cells [11]. We also found that MG2 subgroup expressed genes involved in response to oxidative stress (*SOD2*, *NFKBIA*, *NFKBIZ*, *TNFAIP3*, *BIRC3* and *CD44*). Additionally, MG3 cells expressed several rod markers, *RHO*, *PDE6G* and *GNGT1,* along with traditional Müller cell markers. The proportion of MG3 cells expanded slightly within the peripheral retinal samples after exposure to light stress. This finding is consistent with the previous reports that Müller cells have stem cell characteristics that may help regenerate injured or stressed retina [4, 12]. We also found that, of all the Müller cell subtypes, MG1 cells had the lowest likelihood of light stress-associated relative values (**Figure 3G**), indicating that this subset was relatively unresponsive to light stress. MG2 cells had higher values of light stress-associated relative likelihood, suggesting that this population is more responsive to light stress (**Figure 3H**, red, highly changed).

We performed velocity and trajectory analysis to investigate the transitional relationship between Müller cells and rods. Individual cell velocities were projected onto the UMAP embedding, which was superimposed with Seurat-defined Müller glia subtypes and rods (**Figure 3I**). The Python package PAGA was employed to determine velocity-inferred directionality [13]. We observed a differential trajectory between Müller glia and rods. MG3 exhibited the highest root-like potential and a fast pace of differentiation (**Figure 3I**). This pace was maintained during the production of MG2 cells with high light stress-associated relative likelihood values, which substantially slowed down as the cells became more rod-like (**Figure 3J &K, Supplementary Figure 5G**). These findings indicate a transition trajectory that starts with control-like MG3 cells, followed by light-affected MG3 cells that ultimately result in three different endpoints (MG1, MG2 cells and rods). Remarkably, the light stress associated likelihood increases in correlation with the differential trajectory observed within these three endpoints. However, MG2 cells displayed the most rapid transition pace.

### Target genes involved in the response to light stress in the periphery-dominant Müller cells

Given the significantly different responses to light stress observed between MG1 and MG2 cells, we investigated the top target genes that were region-specific and associated with pathological processes. We selected the top ten genes that exhibited differential expression between MG1 and MG2 with or without light stress (**Figure 4A**). Substantial transcriptomic changes were observed in MG2 cells in response to light stress, in contrast to MG1 cells, which had minimal changes. Among the top ten genes highly enriched in MG2, *MT1F*, *AKAP12*, *MT1X*, *CCN1*, *MT1E*, *MAFF*, *MT1G* were upregulated in response to light stress. These findings indicate that Müller cells in the human retina consist of distinct subtypes with unique transcriptome profiles exhibiting differential stress responses. We chose three top DEGs, *MT1G*, *AKAP12* and *MAFF*, which were particularly abundant in MG2 (**Figure 4B**) and were highly responsive to light stress (**Figure 4C**, highly changed) for further study.

**Figure 4.**
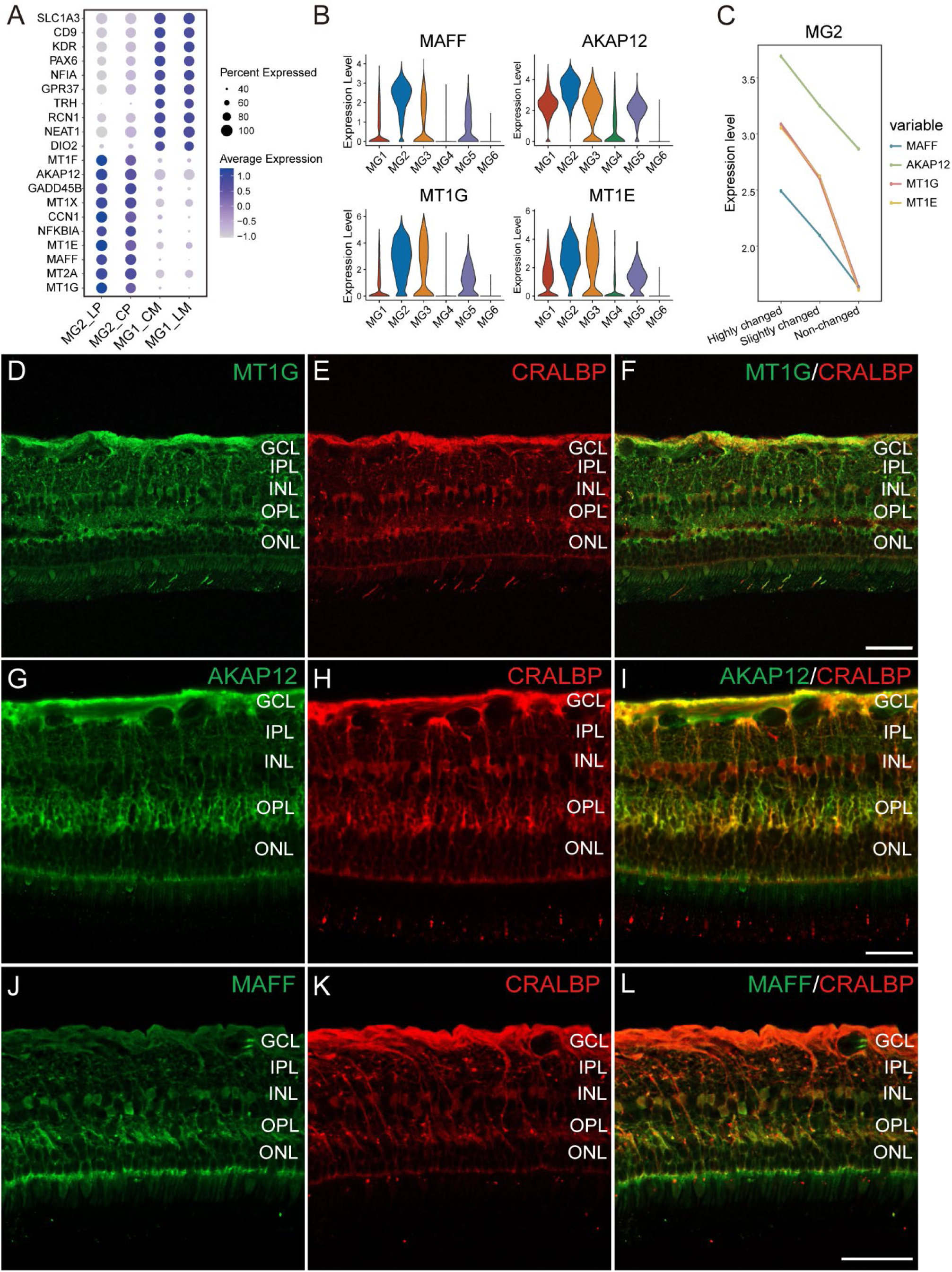
Genes involved in the response to light stress in periphery-dominant Müller cells and immunofluorescent (IF) staining of MT1G, AKAP12 and MAFF in human peripheral retina. **A**. Dot plot illustrating the top ten DEGs between MG1 and MG2 cells in response to light stress. **B**. Violin plot showing the expression levels of *MAFF, AKAP12, MT1G* and *MT1E* in subtypes of Müller cells. **C**. Line chart showing the average expression levels of *MT1G*, *MT1E*, *AKAP12* and *MAFF* in MG2 cells in response to light stress. **D**. IF staining of MT1G (green) in human peripheral retina. **E**. IF staining of CRALBP (red), a Müller cell marker. **F**. Co-localization of MT1G and CRALBP. **G**. IF staining of AKAP12 (green) in human peripheral retina. **H**. IF staining of CRALBP (red). **I**. Co-localization of AKAP12 and CRALBP. **J**. IF staining of MAFF (green) in human peripheral retina. **K**. IF staining of CRALBP (red). **L**. Co-localization of MAFF and CRALBP. Scale bar=50 µM. GCL: Ganglion Cell Layer; IPL: Inner Plexiform Layer; INL: Inner Nuclear Layer; OPL: Outer Plexiform Layer; ONL: Outer Nuclear Layer.

Protein expression patterns of the three top DEGs, *MT1G*, *AKAP12* and *MAFF*, in the human retina, were studied with immunofluorescent staining. We found that MT1G primarily co-localised with cellular retinaldehyde-binding protein (CRALBP), a Müller cell-specific marker (**Figure 4D-F**). AKAP12 staining co-localised with CRALBP in the retinal nerve fibre layer, outer plexiform layer and outer limiting membrane of the peripheral retina (**Figure 4G-I**). We observed MAFF staining in cell bodies of Müller cell and their process in outer plexiform layer and outer limiting membrane. (**Figure 4J-L**). We validated the protein expression of *MT1G*, *AKAP12* and *MAFF* using freshly obtained retinal tissue punches collected from different locations across the human retina (**Supplementary Figure 6**). The protein expression of these genes was consistent with the distinct expression patterns identified in our RNAseq analysis, with lower expression levels observed in the macula than that in the peripheral retina.

### Functional validation of *MT1G*, *AKAP12* and *MAFF* in Müller cells in response to light Stress

We knocked down *MT1G*, *AKAP12* and *MAFF* in primary human Müller cells with siRNA and exposed them to light stress (**Figure 5A**). Knockdown of *MAFF* reduced cell viability by 5-10% compared with control siRNA treated cells under dim light (**Figure 5B**). Under light stress, by contrast, the viability of Müller in which *MT1G*, *AKAP12* or *MAFF* had been knocked down decreased by 10-30% compared with the control (**Figure 5C**).

**Figure 5.**
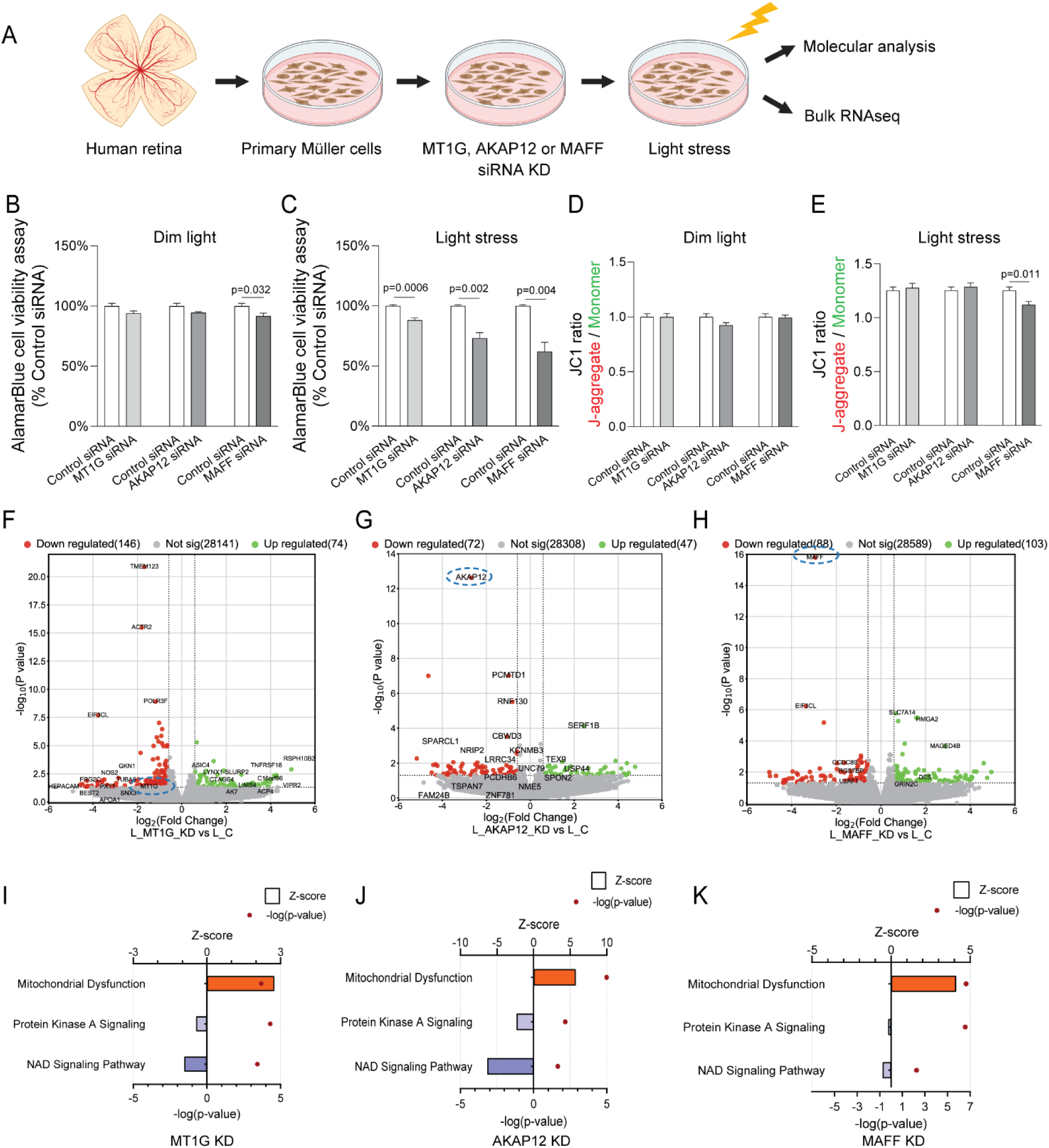
Knockdown of *MT1G*, *AKAP12* and *MAFF* in human primary Müller cells. **A.** *MT1G*, *AKAP12* and *MAFF* siRNA knockdown in primary Müller cells exposed to light stress. **B** & **C**. AlamarBlue cell viability assay on primary Müller cells with or without *MT1G*, *AKAP12* and *MAFF* siRNA treatment in response to light stress. n=6 biological replicates per group. Statistical analysis was performed using Welch’s t-test (two-sided) between the control group (used consistently across all pairwise comparisons) and the respective knockdown groups. Data are presented as means ± standard error of the mean (SEM). **D** & **E**. JC1 assay on primary Müller cells with or without *MT1G*, *AKAP12* and *MAFF* siRNA knockdown in response to light stress. n=8 biological replicates per group. Statistical analysis was performed using Welch’s t-test (two-sided) between the control group (used consistently across all pairwise comparisons) and the respective knockdown groups. **F**-**H**. Volcano plots of differential gene expression (p < 0.05, FC > 1.5) in *MT1G*, *AKAP12* and *MAFF* siRNA knockdown vs. control groups. **I**-**K**. IPA of differential gene expression in human primary Müller cells with *MT1G*, *AKAP12* and *MAFF* siRNA knockdown compared to the control group in response to light stress.

We also evaluated the mitochondrial membrane potential of primary Müller cells using JC-1 staining. The JC-1 ratio (J-aggregate/Monomer) was not significantly different between the control and *MT1G*, *AKAP12* and *MAFF* knockdown groups under dim light (**Figure 5D**). However, the JC- 1 ratio of the *MAFF* knockdown group was significantly lower than that of the control group under light stress (**Figure 5E**). These findings suggest that *MAFF* may play an important role in the Müller cell stress response.

To gain further insights into the function of these genes, we performed bulk RNA sequencing in Müller cells with *MT1G*, *AKAP12* and *MAFF* siRNA treatment, with or without light stress. The mRNA levels of *AKAP12*, *MAFF* and *MT1G* were significantly reduced, as shown in the volcano plots (**Figure 5F-H**, blue circles). Heatmaps of gene expressions between *AKAP12*, *MAFF*, *MT1G* siRNA-treatment groups and the control group displayed distinct clustering of individual samples within the same group (**Supplementary Figure 7**).

Ingenuity Pathway Analysis (IPA) analysis revealed several significant differences in canonical pathways that were impacted by *MT1G*, *AKAP12* or *MAFF* knockdown. The commonly altered pathways included mitochondrial dysfunction, protein kinase A signalling and NAD signalling (**Figure 5I-K**). The transcription level changes suggested that pathways associated with mitochondrial dysfunction were activated in all three target gene knockdown groups, while pathways related to protein kinase A and NAD signalling pathways could have been inhibited.

### Dysregulation of MT1G, AKAP12 and MAFF in Human Diseased Retinas

We analysed the expression of MT1G, AKAP12 and MAFF in postmortem human retinas with documented age-related macular degeneration (AMD) with geographic atrophy and diabetic retinopathy (DR) (**Supplementary Figures 8A-D**). Geographic atrophy in the retinas with AMD were characterised by loss of the photoreceptors and retinal pigment epithelium with extensive Müller cell filaments (fibrotic tissue) filling the space (**Figure 6B, 6F** and **6J**).

**Figure 6.**
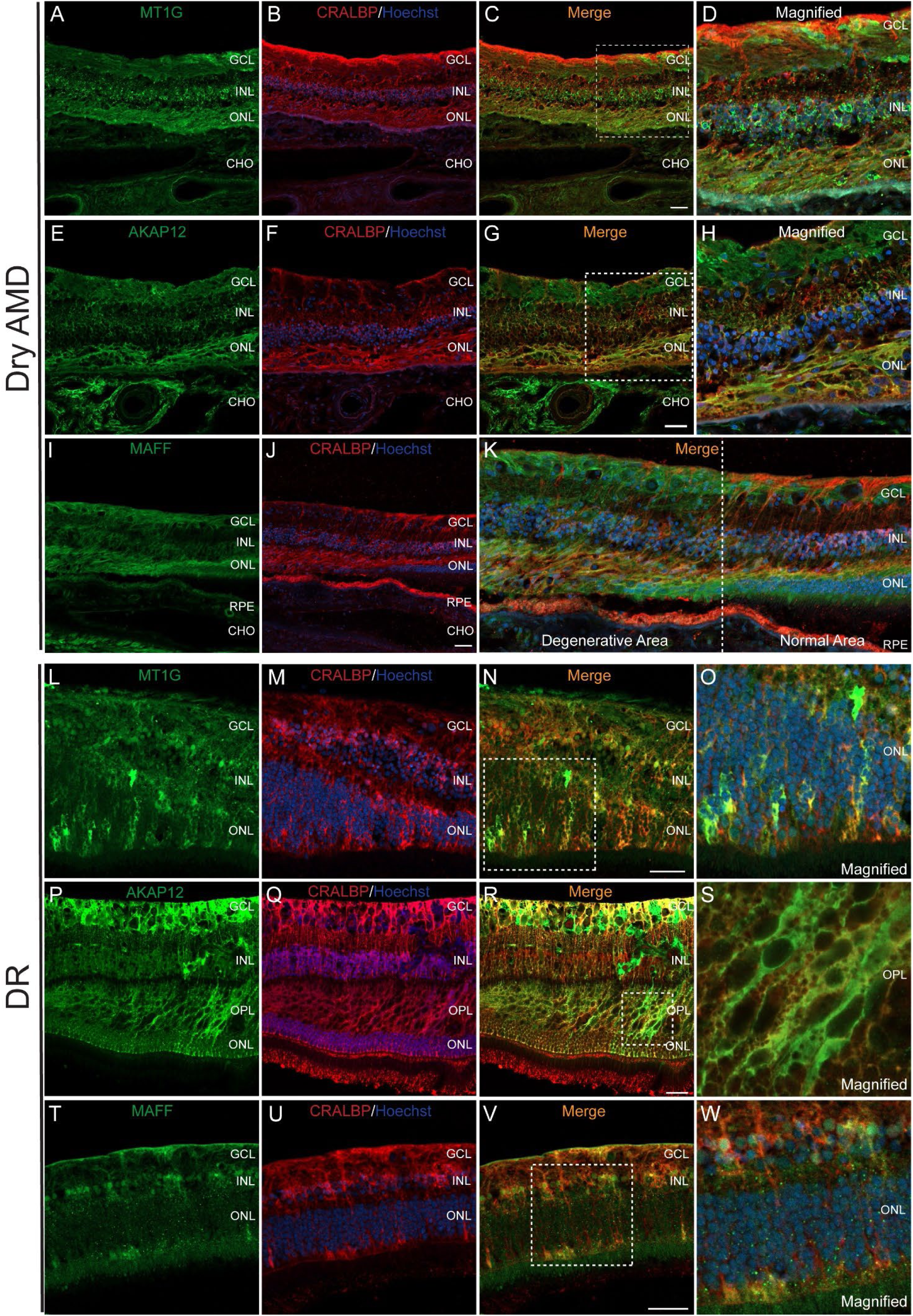
Dysregulation of *MT1G, AKAP12 and MAFF* in human retinas with dry AMD and DR. **A-D**. IF staining of MT1G (green) and CRALBP (red) on the donor retina with dry AMD. **A**. IF staining of MT1G. **B**. IF staining of CRALBP and Hoechst (blue). **C.** Co-localization of MT1G and CRALBP. **D**. Magnified image of the dotted box in **C**. **E-H**. IF staining of AKAP12 (green) and CRALBP (red) on the donor retina with dry AMD. **E**. IF staining of AKAP12. **F**. IF staining of CRALBP and Hoechst (blue). **G**. Co-localization of AKAP12 and CRALBP. **H**. Magnified image of the dotted box in **G**. **I-K**. IF staining of MAFF (green) and CRALBP (red) on the donor retina with dry AMD. **I**. IF staining of MAFF. **J**. IF staining of CRALBP and Hoechst (blue). **K**. Co- localization of MAFF and CRALBP. **L-O**. IF staining of MT1G (green) and CRALBP (red) on the donor retina with DR. **L**. IF staining of MT1G. **M**. IF staining of CRALBP and Hoechst (blue). **N**. Co-localization of MT1G and CRALBP. **O**. Magnified image of the dotted box in **N**. **P-S**. IF staining of AKAP12 (green) and CRALBP (red) on the donor retina with DR. **P**. IF staining of AKAP12. **Q**. IF staining of CRALBP and Hoechst (blue). **R**. Co-localization of AKAP12 and CRALBP. **S**. Magnified image of the dotted box in **R**. **T-W**. IF staining of MAFF (green) and CRALBP (red) on the donor retina with DR. **T**. IF staining of MAFF. **U**. IF staining of CRALBP and Hoechst (blue). **V**. Co-localization of MAFF and CRALBP. **W**. Magnified image of the dotted box in **V**. Scale bar = 50 µM.

We observed abnormal expression of MT1G, AKAP12 and MAFF (green, **Figure 6A, 6E** and **6I**) in these degenerative areas. MT1G staining was still primarily co-localized with the Müller cell marker CRALBP (**Figure 6B-C**). Notably, MT1G exhibited unusually strong staining in the outer nuclear layer (ONL), particularly in the areas of geographic atrophy in the retina with AMD (**Figure 6D**). AKAP12 was still mainly expressed within Müller cells, as evidenced by its co-localization with CRALBP staining (**Figure 6E-H**), with unexpectedly strong staining in the ONL. The immunostaining of MAFF also showed abnormalities. Although MAFF was still co-localized with CRALBP, it appeared upregulated in the degenerated areas, especially at the margins (**Figure 6K**, left). In contrast, the expression level was relatively low in more regular areas (**Figure 6K**, right).

The retina with DR exhibited significant disruption (**Figure 6M** and **6Q**) or thinning (**Figure 6U**) of the inner nuclear layer (INL). We observed unusual, patchy upregulation of all three target proteins—MT1G, AKAP12 and MAFF—within the Müller cells of the DR retina (**Figure 6L, 6N, 6P, 6R, 6T** and **6V**). The patchy activation of MT1G was primarily localised to the ONL (**Figure 6O)**. AKAP12 was patchily activated in the outer plexiform layer (**Figure 6S**). MT1G and AKAP12 were also elevated in the walls of intraretinal cysts (**Supplementary Figures 8E-H**), though this may be partly due to condensation of Müller cells in these regions. The patchy upregulation of MAFF staining was predominantly observed in the cell bodies of Müller cells within the INL and processes in the ONL (**Figure 6W**).

### Dysregulation of MT1G, AKAP12 and MAFF in retinal disease mouse model

We examined the protein expression of *MT1G, AKAP12* and *MAFF* in JR5558 mice that spontaneously develop bilateral subretinal neovascularisation [14, 15]. Subretinal lesions appeared around 4 weeks and became established at 8 weeks, as confirmed by colour fundus photography, optical coherence tomography and fundus fluorescein angiography (**Figure 7A-C**). We compared the protein expression of *MT1G, AKAP12* and *MAFF* in the JR5558 mouse retina with the C57BL/6 control retina. We found that AKAP12 was upregulated at both 4 weeks and 8 weeks of age in the JR5558 mouse retina compared with the control retina (129% vs 100%, 169% vs 100%) (**Figure 7 D&E**). The protein expression of *MAFF* and *MT1G* was significantly greater at 8 weeks of age in the JR5558 mouse retina than in the control (217% vs 100%, 121% vs 100%, respectively). These results show that the stress-responsive genes *MT1G, AKAP12* and *MAFF* are activated in subretinal neovascularisation, which is a common vision-threatening mechanism in retinal disease.

**Figure 7.**
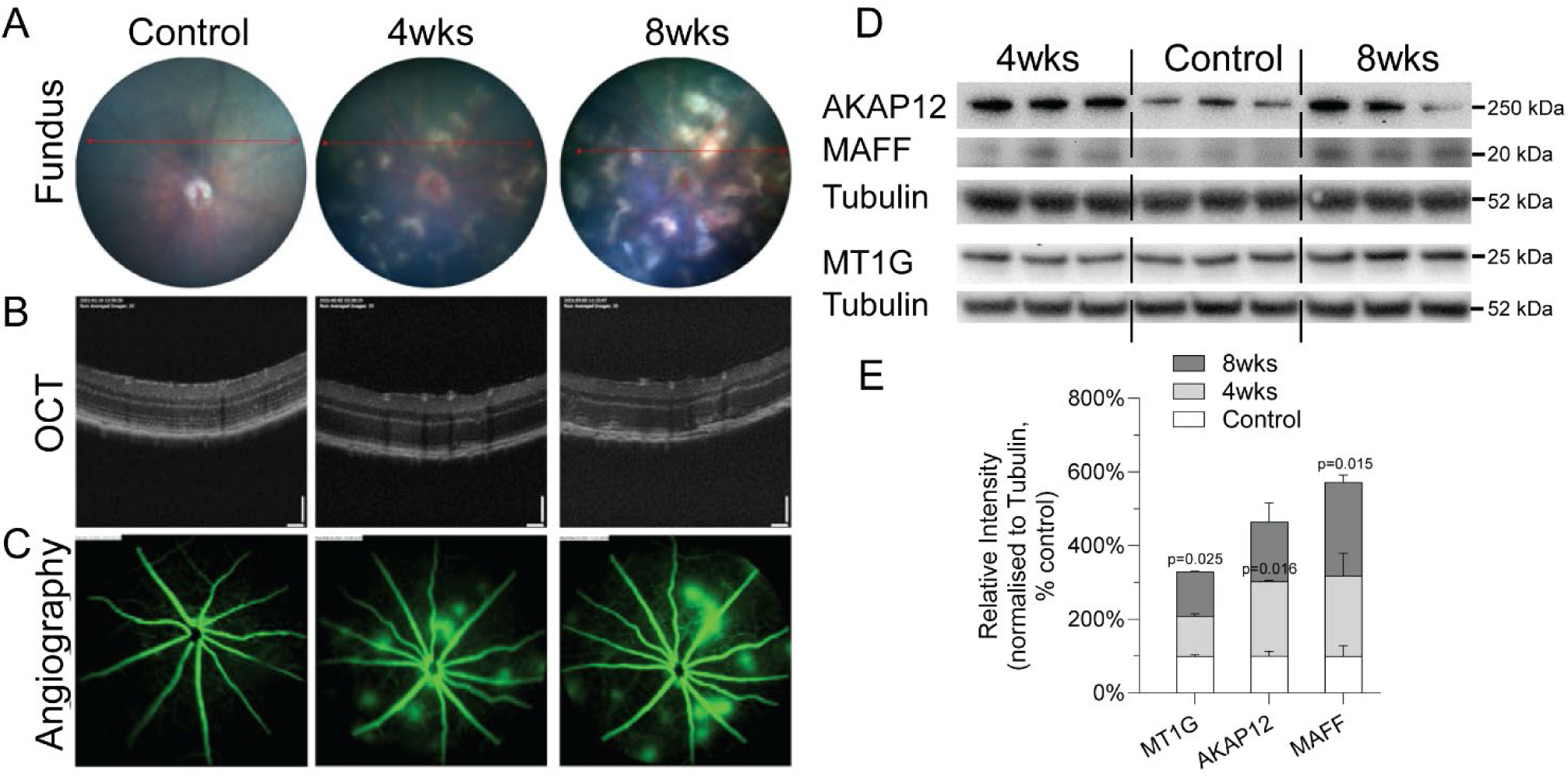
Dysregulation of *MT1G, AKAP12* and *MAFF* in JR55558 mice mimicking subretinal neovascularisation. **A**. fundus photos (**A**), OCT (**B**) and FFA (**C**) images of the C57BL/6J control and JR5558 mice at 4 and 8 weeks of age. **D & E**. Images of representative protein bands and the corresponding densitometry analysis of Western Blot against MT1G, AKAP12 and MAFF in the retinas of both control and JR5558 mice at the ages of 4 and 8 weeks of age. n=3 mouse retinas per group. Statistical analysis was performed using Welch’s t-test (two-sided) between the control and the 4 weeks or 8 weeks old groups, respectively.

## Discussion

Though Müller cells are found in both the macula and the peripheral retina, disparities exist in their distribution, density, morphology and molecular functions across these regions [5]. This study examined the stress responses of Müller cells in both areas utilising scRNAseq on human macular and peripheral retinal explants. Macular Müller cells had minimal transcriptional changes in response to stress compared with their peripheral counterparts. Analysis of the top three genes most abundantly expressed in peripheral Müller cells revealed a crucial role in cellular survival under stress. Our findings indicate a previously unobserved distinction between macular and peripheral Müller cells, which might offer insights into the vulnerability of the macula in specific retinal diseases such as AMD, macular oedema and MacTel.

Metallothionein (*MT*) refers to a group of low molecular weight, cysteine-rich proteins that play critical roles in various biological processes, including metal metabolism [16], protection against DNA damage [17] and regulation of apoptosis [18, 19]. *MT1* has been implicated in age-related neurodegeneration [20, 21], as it is associated with aging and endoplasmic reticulum (ER) stress, particularly in defending against nitric oxide (NO)-induced stress. NO-induced stress in Müller cells has been implicated in the development of various retinal conditions, such as DR, glaucoma and AMD [22].

*AKAP12* interacts with a variety of key kinases, such as PKA and PKC, which are essential for the proper regulation of morphogenetic cell movement [23]. *AKAP12* is involved in maintaining vascular integrity and regulating endothelial cell function [24–26]. It has been identified as a maturation factor for the blood-brain barrier (BBB) that alleviates damage and dysfunction of the BBB caused by ischemic stress [27]. Although the specific function of *AKAP12* in human Müller cells remains largely unclear, the increased expression in Müller cell processes within the ONL of AMD eyes, along with the patchy upregulation observed, particularly around areas with evident intraretinal fluid in DR retinas, suggests a potential link to the response of Müller cells to retinal pathology.

*MAFF* protein belongs to the small Maf family of proteins that are known to form heterodimers with Nrf2, thereby modulating the transcriptional activity of downstream genes [28, 29]. There is limited information regarding the specific role of *MAFF* proteins in human diseases, but they have been implicated in various pathological conditions such as cancer, diabetes, immune system disorders and neurodegenerative diseases such as Alzheimer’s disease and Parkinson’s disease [30–34]. The role of *MAFF* in Müller cells has not been previously investigated. However, its elevated expression in Müller cells, especially at the margins around the degeneration area in the AMD, suggests it may modulate the Nrf2-related stress response in these cells. Further study is warranted to elucidate the roles of MAFF in Müller cells and other biological processes, this is beyond the scope of the current study.

*MT1G*, *AKAP12* and *MAFF* are much more strongly expressed in Müller cells of the peripheral retina than of the macula. siRNA knockdown of these genes in human primary Müller cells disrupted multiple stress response signalling pathways, including mitochondrial function and the protein kinase A and NAD signalling pathways. A better understanding of how the macula responds to stress when it only expresses stress response genes weakly and non-inducibly may pinpoint new treatment targets for macular diseases.

Macular Müller cells have a distinct phenotype, with a weaker expression of stress response genes than Müller cells from the peripheral retina. This unique phenotype may offer an evolutionary advantage by preserving the normal function of the macula under stress, thereby maintaining central vision during sudden environmental changes and increasing survival chances in harsh conditions. Cellular stress responses typically promote cell survival, but may also compromise normal cellular functions [35]. An acute stress response in Müller cells might disrupt their ability to effectively regulate neurotransmitter levels, leading to an imbalance in the retinal chemical environment and short-term vision impairment. Through minimising their stress response, macular Müller cells may have evolved to suppress their stress response in favour of maintaining their normal function under acute stress to preserve central vision in a rapidly changing environment. This adaptation would have been beneficial in evolution when central vision was critical for as hunting, foraging and evading predators. The capacity to maintain clear central vision during abrupt changes in lighting condition would have substantially enhanced their chances of survival in harsh conditions.

Studying the macula is challenging because laboratory animals do not have maculas apart from non-human primates which raise ethical issues. We addressed this by using *postmortem* human retinal/macular explants to investigate the stress response of the human macula. The data derived from this model system complement other models utilised in retina research, such as mice and organoids. One of the primary advantages of using human retinal explants is the direct examination of gene expression and morphology of the human macula in its entirety. Inherent limitations of this system include the lack of a functional vascular network and interaction with the retinal pigment epithelium (RPE). These disadvantages could potentially be addressed with perfusion and co-culture systems. The use of postmortem tissue may raise concerns regarding viability and metabolic activity, but there is growing evidence to support the functionality of cultured human retinal explants despite *postmortem* delay [36]. We have previously reported that the metabolic features of human retinal explants at least as good as those of mouse retinas and retinal organoids [37].

Our use of acute intense light exposure as a model for the chronic stress associated with aging may not perfectly reflect the true *in vivo* response of the human macula and peripheral retina. Despite this, our method is a valuable starting point for investigating the stress response of the macula. It sets the stage for future, in-depth research on neurotoxins, such as deoxy-sphinganine (unpublished data), that significantly influence the pathogenesis of macular telangiectasia type 2 [38] and it helps us begin to unravel the distinct ways the macula responds to stress.

Exposure to light stress induced distinct patterns of cellular responses in the peripheral retina and the macula. Our single-cell RNA sequencing analysis revealed that both Müller cells and rods exhibited significant differential gene expression in the peripheral retina. Cones, horizontal cells and bipolar cells also showed differential gene expression, though to a lesser extent. We focused on the Müller cell response to light stress primarily because Müller cells presented two subpopulations with distinct transcriptomic profiles—one predominantly in the macula and the other in the peripheral retina. Müller cells are among the first responders to retinal stress, playing crucial roles in maintaining retinal homeostasis and responding to injury [3]. Additionally, it is well established that the phototransduction cascade in photoreceptors is one primary source of oxidative stress and cell death [9]. Müller glial cells also play a crucial role in the cone Müller visual cycle, a specialized pathway that supports the rapid regeneration of visual pigments in cone photoreceptors, which are essential for colour vision and high-acuity vision in bright light [39, 40]. Therefore, the transcriptomic differences observed in Müller cells between the periphery and the macula may also be influenced by the varying densities of rods/cones in these regions, with a lower proportion of rods present in the macula.

Amacrine cells, which play a crucial role in modulating signals between bipolar and ganglion cells, may have evolved to meet the high visual demands of the macula region. This specialization could make them more sensitive to changes in light conditions, ensuring accurate and rapid visual perception. Although the alteration score for amacrine cells is highest in the macula, the number of amacrine cells detected in our dataset is relatively low (1,071 out of 77,405 cells, ∼1.4%). This limitation is a recognized challenge of the scRNAseq technique, where there is less confidence in the findings from cell subpopulations that are only present in small numbers. Consequently, while we acknowledge the potential importance of the observed changes in amacrine cells, the current dataset does not provide sufficient resolution to draw definitive conclusions.

In conclusion, we observed that macular Müller cells exhibit a subdued response to stress, possibly due to the higher functional demands placed on them in the macula region. These cells express low levels of several key stress-response genes that significantly contribute to the resilience of peripheral Müller cells under stress. Future investigations are needed to clarify further the molecular mechanisms underlying the distinct stress response characteristics of macular Müller cells. Our findings indicate that this reduced stress response in macular Müller cells could offer an evolutionary advantage by allowing these cells to sustain their normal functions even under acute stress.

## Materials and Methods

### Tissue Collection and Retinal Dissection

The use of donated human eyes was approved by the Human Research Ethics Committee of the University of Sydney (Protocol Number: 2016/282 and X21-0208 & 2020/ETH03339). Donors had no known eye diseases. Retinal dissection was conducted as previously described [41]. In brief, donor eyes were acquired from the New South Wales Tissue Bank after the corneas had been removed for transplantation. The donor issues were preserved in CO_2_-independent medium at 4 °C until they were dissected. For dissection, the eye was placed on a petri dish and the iris, lens and vitreous were excised from the eyecup. The neural retina was then carefully detached from the underlying retinal pigment epithelium and transferred to a fresh petri dish. Circular retinal tissues with a diameter of 5mm were collected from the macula or mid-peripheral region of the superior retina by punch biopsy (#BP-50F, Kai Medical) and processed for downstream assays. The mid-peripheral region was identified as the midpoint between the fovea and ora serrata. 2mm-diameter circular retinal samples extending from the fovea to the superior and inferior peripheral neural retina (**Supplementary Figure 6A**) were collected and extracted for protein analysis. One postmortem eye from a donor with diabetic retinopathy (DR) was used to validate the expression of target genes in the diseased retina. Detailed fundus photography was conducted to document characteristics and features present within the diseased retina.

### Light and High Glucose Stress Models on Retinal Explants and Human Primary Müller Cells

We employed a custom-built light exposure system situated within the tissue culture incubator to induce light stress in retinal explants and human primary Müller cells (**Supplementary Figure 1**). Light stress was accomplished by subjecting explants to intense light at 32k lux, while the control group were exposed to 5 lux light for a continuous period of 4 hours. The light source was positioned at a distance of 10 cm from the samples in both the experimental and control groups to ensure uniform light exposure. Retinal explant tissues were placed on 12-well plate transwell inserts (#3460, Corning) with neurobasal medium (#21103049, Gibco), 1% B27 (#17504044, Gibco), 1% N2 (#17502048, Gibco), 1% fetal bovine serum (FBS, #F9423, Sigma-Aldrich), 1% GlutaMax (#35050, Gibco), 1%ITS (#51500056, Gibco) and 1% penicillin-streptomycin (P/S, #P4333, Sigma-Aldrich). Macular and peripheral neural retinal explants were exposed to light for 4 hr (Fig1A). Three days after siRNA transfection, human primary Müller cells in 12-well (#3513, Corning) or 96-well plates (#3340, Corning) were exposed to intense (32k lux) or dim (5 lux) light. Macular and peripheral retinal explants were cultured in either normal glucose (5mM glucose, left eye) or high glucose (25mM, right eye of the same donor) on 12-well transwell inserts with DMEM (#10569, Gibco) supplemented with 1% B27, 1% N2, 1% FBS and 1% P/S for 24 hours.

### Single Cell dissociation and cDNA library construction

Macular and peripheral explants were dissociated into single cells following light or high glucose treatment as depicted in **Figure 1A**. Single-cell dissociation was achieved using the Papain Dissociation System (Worthington Biochem, NJ, USA) according to the manufacturer’s protocol. Briefly, the entire retinal explant was immersed in papain solution for 30 minutes at 37 °C, followed by dissociation through pipetting with a glass pipette. The single-cell suspension was combined with barcoded gel beads in a microfluidic chip and run on the Chromium Controller (10X Genomics) to create gel bead-in-emulsion (GEMs) droplets (Chromium Next GEM Single Cell 3ʹ Kit v3.1, 16 rxns PN-1000268, 10×Genomics). The GEMs were incubated, enabling mRNA reverse transcription into cDNA. The barcodes from the gel beads were incorporated into the cDNA molecules by uniquely tagging the cDNA originating from each individual cell. The barcoded cDNA was then amplified using PCR to construct a library. The libraries were sequenced on an Illumina NovaSeq 6000 platform.

### Single Cell RNA sequencing data analysis

The raw sequencing data was processed by the Cell Ranger software (v3.0.2, 10x Genomics). A human reference genome (GRCh38) was used to accurately map the sequencing readouts and quantify gene expression. We then applied Seurat (v4.1.1) [42] to perform the pre-processing and downstream analysis of the single-cell gene expression data. Cells with an expression of unique feature counts over 6,000 (as for the high glucose treated group, this parameter was 10,000) or less than 200, a mitochondrial gene ratio over 15% and a hemoglobin gene ratio over 5% were filtered out. DoubletFinder [43] (v2.0.3) was used to detect and remove the doublets as to the following criteria: pN = 0.25, pK = 0.09, PCs = 1:20, sct = TRUE and reuse.pANN = FALSE. After the above quality control, we normalised and log-transformed the featured expression measurements for each cell. 2,000 highly variable features were chosen for downstream data scaling. The integration of multiple single-cell datasets from different biological contexts was implemented by SeuratWrappers (v 0.3.0) with LIGER algorithm. It depends on integrative non-negative matrix factorisation to delineate shared and dataset-specific factors of cell identity [44]. Then the “FindNeighbors” and “FindClusters” functions were applied for unsupervised clustering. For visualization, UMAP (Uniform Manifold Approximation and Projection) was used to reduce the dimensionality. We further annotated each cell cluster based on specific marker gene expression. The differentially expressed genes were calculated by the “FindMarkers” function with specific settings (only.pos=FALSE, min.pct=0.2, logfc.threshold = 0.2)

### Calculating the Alternation Score of different cell types

To quantify the change in cellular response to light, we designed the Alternation (AC) Score. The first step of selecting the signature genes was conducted as a previously published method that characterized the MI contribution [10]. However, we have modified the second step to calculate both the quantity and expression level change of signature genes during the light stress process. Here, we defined the new *AC_score_* as follows:

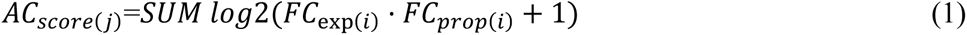

In which *FC_exp_*_(*i*)_ refers to the expression fold change of the i^th^ signature gene in the j^th^ cell type between the light stress and control groups, *FC_prop_*_(*i*)_ is the proportion fold change of the cells that express the i^th^ signature gene between the light stress and control groups in the j^th^ cell type.

### Light stress-associated relative likelihood analysis

MELD algorithm implemented in Python (v.3.8.16) was employed to calculate the continuous light stress-associated relative likelihood of each cell across the entire single-cell dataset [45]. In brief, the density of each sample (replicates and conditions) was estimated on a MELD graph built over the cellular transcriptomic state space (beta=67, knn=7). We then compared the density estimates between strong and dim light conditions within each replicate. The resulting relative likelihoods across replicates were normalized to quantify the probabilities of a given cell would be observed in the light stress condition. Cut-offs of values less than 0.4, between 0.4 and 0.6 and greater than 0.6 were used to determine whether a cell was non-changed, slightly changed, or highly changed, respectively.

### Single-cell Trajectory Analysis

We utilized the 10×velocyto pipeline on the filtered cell ranger-generated BAM files to determine the number of spliced and unspliced reads for each sample. Then, we employed the dynamical model of scVelo v.0.2.3 [46] for single-cell RNA velocity inference. We obtained a comprehensive view of RNA velocity across all genes and cells by visualizing a vector field on top of the 2D dimensional reduction plot. We enhanced our interpretation by leveraging partition-based graph abstractions (PAGA) to ascertain the directionality between cells.

### Gene Ontology Analysis

Gene Ontology (GO) analysis was conducted using the web-based resource, the Generic Gene Ontology Term Finder [47], accessible at https://go.princeton.edu/cgi-bin/GOTermFinder. Highly expressed genes in macula-dominant Müller glia (m-dMG) were defined as “m-HXs”. Highly expressed genes in peripheral retina-dominant Müller glia (p-dMG) were defined as “p-HXs”. Light-induced Changes were defined as “LiC”. GO analyses were performed on the m-HXs and p-HXs (fold change >1.5,) with (adjusted p value <0.05) or without LiC (adjusted p value >0.05).

### Ingenuity Pathways Analysis

QIAGEN Ingenuity Pathways Analysis (IPA) was performed to identify significant canonical pathways and predict which pathways are activated or inhibited between control and MT1G, MAFF and AKPAP12 siRNA treatment groups. Upregulated and downregulated genes were imported into the IPA software and subjected to core analysis to identify canonical pathways.

### Vibratome sectioning, immunofluorescent staining and imaging

Human retinal tissue from the macula and mid-periphery was obtained using a 5mm-diameter biopsy punch after separating the neural retina from the RPE-choroid-sclera eyecup. Tissues were fixed in 4% paraformaldehyde for 1 hour and then washed with 1× phosphate-buffered saline (PBS). Retinal tissues were embedded in 3% low-melting agarose (#50100, Lonza) within a plastic cubic mould. The tissue blocks were secured onto the specimen plate of a vibratome (#VT1200S, Leica). Serial tissue sections, 100µm thick, were generated through vibrating sectioning. The tissue slices were maintained in cold 1× PBS until further use. Retinal slices were blocked overnight in blocking buffer (5% normal donkey serum and 0.5% Triton-X100 in PBS) in a 48-well plate (#3548, Corning). Primary antibodies (**Table 1**) were diluted in 1% normal donkey serum and 0.5% Triton-X100 in PBS. Tissues were incubated in primary antibody for consecutive 5 days, then washed three times with PBS and placed in a diluted secondary antibody (1% normal donkey serum and 1% Triton-X100 in PBS) for two days. Tissues were then washed 3 times with PBS. Nuclei were stained with Hoechst 33342 for 30 minutes in the dark. Slices were mounted onto polysine-coated slides (Menzel Glaser, Germany) with secure-seal spacers (ThermoFisher Scientific, MA, USA) using VECTASHIELD mounting medium (#H-1000, Vector Laboratories). Confocal images were taken using a confocal microscope and imaging system (Zeiss LSM700). Zen Software (blue edition) from Zeiss was used to analyse the images.

**Table 1.** List of Antibodies Used in the Study.

| Antibodies | Source | Catalogue | Host | Dilutions |  |
| --- | --- | --- | --- | --- | --- |
|  |  | NO. |  | WB | IF |
| $\alpha/\beta$-Tubulin | Cell signalling | #2148 | Rabbit | 1:1000 | N/A |
| CRALBP | Abcam | AB15051 | Mouse | N/A | 1:500 |
| MAFF | Abcam | AB183859 | Rabbit | 1:1000 | N/A |
| MAFF | Novus | NBP2-56792 | Rabbit | N/A | 1:200 |
| AKAP12 | Sigma | HPA006344 | Rabbit | 1:1000 | 1:200 |
| MT1G | Abcam | AB193329 | Rabbit | 1:1000 | 1:200 |
| CRALBP | Abcam | AB1243664 | Rabbit | N/A | 1:500 |
| Anti-rabbit-488 | Invitrogen | A21206 | Donkey | N/A | 1:1000 |
| Anti-mouse-594 | Invitrogen | A21203 | Donkey | N/A | 1:1000 |
| Hoechst 33342 | Thermo Fisher | H3569 | N/A | N/A | 1μg/mL |
| Anti-rabbit IgG | Cell signalling | #7074 | Goat | 1:5000 | N/A |
| Anti-mouse IgG | Cell signalling | #7076 | Horse | 1:5000 | N/A |

### Cryosectioning and immunofluorescent staining

The human AMD eyes were fixed in 4% paraformaldehyde (PFA) prepared in PBS buffer for 24 hours. After fixation, the macular retina, choroid and sclera were dissected together and transferred to 20% sucrose in PBS for cryoprotection, incubating overnight at 4°C. The tissue was then embedded in optimal cutting temperature compound and sectioned at a thickness of 16 μm onto Superfrost glass slides. For immunofluorescent staining, tissue sections were blocked in a solution containing 5% donkey serum and 1% Triton X-100 in PBS for 1 hour to prevent non-specific binding. The primary antibody was diluted in PBS containing 1% donkey serum and 1% Triton X-100 and incubated with the sections for 48 hours at 4°C. Following primary antibody incubation, the sections were washed and then incubated with the secondary antibody diluted in PBS containing 1% donkey serum for 4 hours at room temperature. Nuclei were stained with Hoechst33342 for 5 minutes at room temperature. After the staining, the tissue sections were washed three times with PBS and mounted using VECTASHIELD Antifade Mounting Medium (Vector Laboratories, Burlingame, CA, H-1000) before placing coverslips for imaging.

### Western Blot

Samples of 2mm diameter retinal punches were collected from donor eyes (**Supplementary Figure 1A**) and homogenized on ice using radio-immunoprecipitation assay (RIPA) buffer (#900000064244, Merck) supplied with phosphatase and protease inhibitor (1:100, #5872S, Cell Signaling Technologies). Human primary Müller cells were lysed in RIPA buffer with phosphatase and protease inhibitor in a 12-well plate. Tissue and cell lysate were centrifuged at 12,000 g at 4°C for 15 minutes. Supernatants were collected and aliquoted. Protein concentration was determined using a BCA assay kit (#QPBCA, Sigma-Aldrich). Protein samples were mixed with DTT (#D9779-10G, Sigma) and loading sample buffer (#NP0007, Invitrogen), heated at 75°C for 10 minutes and then cooled on ice for two minutes. Samples were loaded onto the 3-8% Tris acetate gel (#EA03785BOX, Invitrogen) for AKAP12 evaluation and electrophoresed at 150 V, 4 °C for 60 minutes. Samples were loaded onto 10-20% Tricine gels (EC66255BOX, Invitrogen) for the evaluation of MAFF and MT1G and were electrophoresed at 125 V, 4 °C for 90 minutes. Proteins were transferred onto PVDF membranes (#IEVH85R, Millipore, USA) using the wet transfer system (#1703930, BioRad, USA), following the manufacturer’s instructions. PVDF membranes were blocked with 5% bovine serum albumin (BSA, #A9647-100G, Sigma) in 1×tris-buffered saline (TBS) for one hour at room temperature and then incubated with primary antibodies (**Table 1**) diluted in 1% BSA in tris-buffered saline with tween 20 (TBST) overnight at 4°C. Membranes were then washed three times with TBST and incubated with secondary horseradish peroxidase (HRP)-antibodies in 1% BSA in TBST for 2 hr. Membranes were then washed three times with 1×TBST and twice with 1×TBS. Protein bands were visualised using Clarity ECL substrate (#170-5061, BioRad) and photographed using the G-Box imaging system (In Vitro Technologies). The specific bands of interest were quantified using the GeneTools image scanning and analysis package. Protein expressions were normalised to α/β tubulin.

### Isolation and culture of primary Müller cells from human donor retinas

Human primary Müller cells were cultured and passaged following the method described previously [41]. Human donor eyes (**Supplementary Table 2**), with corneas removed, were transported in CO_2_- independent medium (#18045088, ThermoFisher Scientific). Retinas were detached from the eyecups, cut into 1 cm² pieces and stored in Dulbecco’s modified Eagle medium (DMEM, #10569010, ThermoFisher Scientific) without FBS or growth factors at 4°C overnight in the dark. The next day, retinal tissue was digested in pre-warmed TrypLE (#12563029, ThermoFisher Scientific) at 37°C for 60 minutes, then transferred to a culture dish with complete medium and cut into ∼1 mm² pieces. Retinal pieces were placed into T25 cell culture flasks (#3289, Corning), separated with an angled 18 G needle and pressed firmly onto the flask bottom to enhance attachment. Flasks were incubated vertically for 15 minutes in the 37°C incubator, then horizontally with 2 ml complete medium (DMEM+10% fetal bovine serum +1% Penicillin-Streptomycin). On day 7, an additional 2 ml of medium was added into the T25 cell culture flask. Routine culture involved medium changes twice a week and after 2-3 weeks, human Müller cell colonies emerged, reaching confluency in another 2-3 weeks. For passaging, cells were rinsed with PBS, digested with TrypLE, centrifuged and resuspended in complete medium, then seeded in new flasks. Subsequent passaging was done at 1:1 initially, progressing to 1:2 or 1:3 ratios after P1, with experiments typically using P3 cells.

### siRNA treatment

Human primary Müller cells (P3) at 80% confluency were transfected with *MT1G* (assay ID: s194623, ThermoFisher Scientific), *AKAP12* (assay ID: s18435, ThermoFisher Scientific), *MAFF* (assay ID: s24370, ThermoFisher Scientific) and control small interfering RNA (siRNA) (assay ID: 4390843, ThermoFisher Scientific) at a concentration of 10nM using Lipofectamine 3000 (#L3000-008, Invitrogen), following the manufacturer’s instructions. Müller cells were treated with siRNA in either 12-well or 96-well plates for three days, reaching 100% confluency prior to further experiments.

### AlamarBlue assay

Human primary Müller cells (P3) were transfected with *AKAP12*, *MT1G*, *MAFF* and control siRNA in a 96-well plate for three days, then starved overnight and exposed to light stress for four hours. Cells were treated with AlamarBlue reagent (1:10 dilution, #DAL1100, ThermoFisher Scientific) and incubated at 37 °C for 4 hr using the AlamarBlue cell viability assay kit. The fluorescence was then read using the microplate reader (Fluostar Omega, BMG Labtech) with the 544nm excitation wavelength and 590nm emission wavelength.

### JC1 assay

Human primary Müller cells (P3) after siRNA treatments were incubated with JC-1 dye (#T3168, Invitrogen) at a concentration of 1.0 μg/mL in DMEM at 37 °C for 30 minutes to measure the mitochondrial membrane potential. The medium was replaced with Hanks’ balanced salt solution (HBSS) and fluorescence was measured using a plate reader. The fluorescence was detected with the 475nm excitation wavelength and 530nm for green emission and 590 nm for red emission. The ratio of red fluorescence divided by green fluorescence of each sample was calculated. All data were normalised to the control siRNA treatment group.

### mRNA extraction, bulk RNA sequencing and bioinformatic analysis

Total mRNAs were extracted from human primary Müller cells treated with *AKAP12*, *MT1G*, *MAFF* and control siRNA using the GenEluteTM Single Cell RNA Purification Kit (#RNB300, Sigma-Aldrich), following the manufacturer’s instructions. The concentration of mRNA was determined using the Qubit RNA HS Assay Kit (#Q32852, Invitrogen). The extracted mRNA samples were sent to Novogene (Singapore) for bulk RNA sequencing after quality control assessment. The library was prepared and a 150 bp paired-end sequencing strategy was used to sequence the samples. The quality of the library was analysed using the Agilent Bioanalyzer 2100 system.

### Mouse eye examinations and retinal tissue collection

Approval for all experimental procedures was obtained from the University of Sydney Animal Ethics Committee, ensuring adherence to ethical standards and the welfare of animals (Project number: 2021/2013). The study utilised JR5558 mice (strain #:005558; RRID:IMSR_JAX:005558) obtained from The Jackson Laboratory for validating the expression of AKAP12, MT1G and MAFF proteins in diseased mouse retinas. C57BL/6J wild-type mice served as the control group. Mice were housed in individually ventilated cages under specific pathogen-free conditions, with autoclaved corn cob bedding and enrichment materials. The environment was controlled at 22 ± 2°C, 50 ± 10% humidity, and a 12-hour light/dark cycle.

JR5558 and C57BL/6J mice were anesthetized using an intraperitoneal injection of ketamine (48 mg/kg) combined with medetomidine (0.6 mg/kg). This ensured the mice were adequately sedated for the imaging procedures. One to two drops of a solution containing 1% tropicamide and 0.5% phenylephrine were administered to each eye to achieve proper pupil dilation, which is critical for high-quality retinal imaging.

Colour fundus imaging was performed using the Phoenix-Micron imaging system (Phoenix MICRON IV) to visualize and document the retinal lesions in the mice. Optical coherence tomography (OCT) was conducted using the Phoenix MICRON OCT2 system, providing detailed transverse images of the retinal layers and structures, which is essential for understanding the extent and nature of retinal lesions.

Fundus fluorescein angiography (FFA) was performed to assess retinal vasculature and perfusion. A 10% fluorescein solution was prepared and administered via intraperitoneal injection. Following the injection, FFA images were captured from both eyes using the Phoenix-Micron imaging system (Phoenix MICRON IV), starting between 30 seconds to 2 minutes post-injection. This allowed for a dynamic assessment of the retinal vasculature, revealing details about blood flow and identifying any abnormalities in retinal perfusion.

The collected retinal tissues at 4 and 8 weeks of age were processed for protein extraction. Western blot analysis was performed to validate the expression levels of AKAP12, MT1G and MAFF proteins.

### Statistical analysis

Each in *vitro* experiment was independently conducted three times from three donors, with more than six replicates per group. The western plot for mouse samples was performed using three independent samples in each group. The data are presented as means ± standard error of mean (SEM). Statistical analysis was performed using Welch’s t-test (two-sided) between the control group (used consistently across all pairwise comparisons) and the respective treatment groups (GraphPad Prism version 10.0.2 for Windows, GraphPad Software, Boston, Massachusetts USA). Assumptions of normality were assessed using the Shapiro-Wilk test. A significance level of p < 0.05 was adopted to indicate statistical significance.

## Data availability

The single-cell RNA sequencing data generated for this study have been deposited in the NCBI Gene Expression Omnibus (GEO) under the accession number GSE249894. The data can be accessed using the secure token ‘qdcdoyigvzydjuj’.

## Declaration of interest

The authors declare that they have no conflict of interest.

## Acknowledgement

All human eye tissues utilized in this research were sourced from the New South Wales Tissue Bank and provided through the Australian Ocular Biobank. This study received financial support from organizations including the Lowy Medical Research Institute (for MG, LZ and TZ), the National Health and Medical Research Council (Investigator Grant GNT_1195021 for MG, Idea grant GNT_ 2020950 for LZ), the Australian Vision Research (for MG, LZ and TZ) and the National Eye Institute (EY032462 and EY031324 for JD). The Innovation Team and Talents Cultivation Program of National Administration of TCM (ZYYCXTD-D-202002, X.F.), the Fundamental Research Funds for the Central Universities (226-2023-00114 and 226-2022-00226, X.F.). The authors thank the High-Performance Computing Cluster of Zhejiang University Innovation Centre of Yangtze River Delta for their technical support.

## Supplementary Figures and Table

**Supplementary Figure 1.**
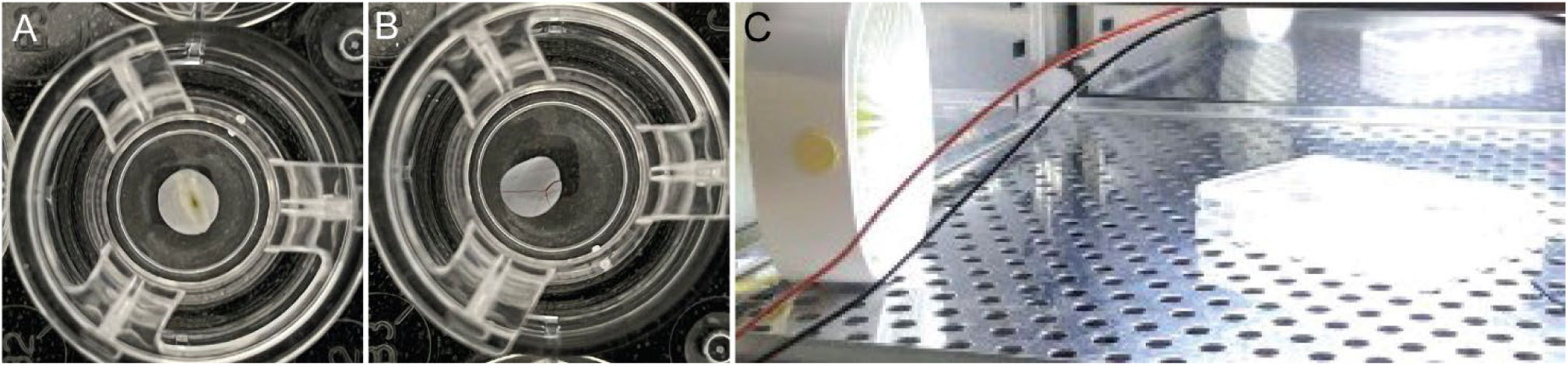
The light stress model of human postmortem macular and peripheral neural explants. The human macular retinal explant (**A**) and mid-peripheral retinal explant (**B**) cultured on transwells. **C**. Light exposure on the retinal explants in transwell plates in a cell culture incubator with temperature control and ventilation.

**Supplementary Figure 2.**
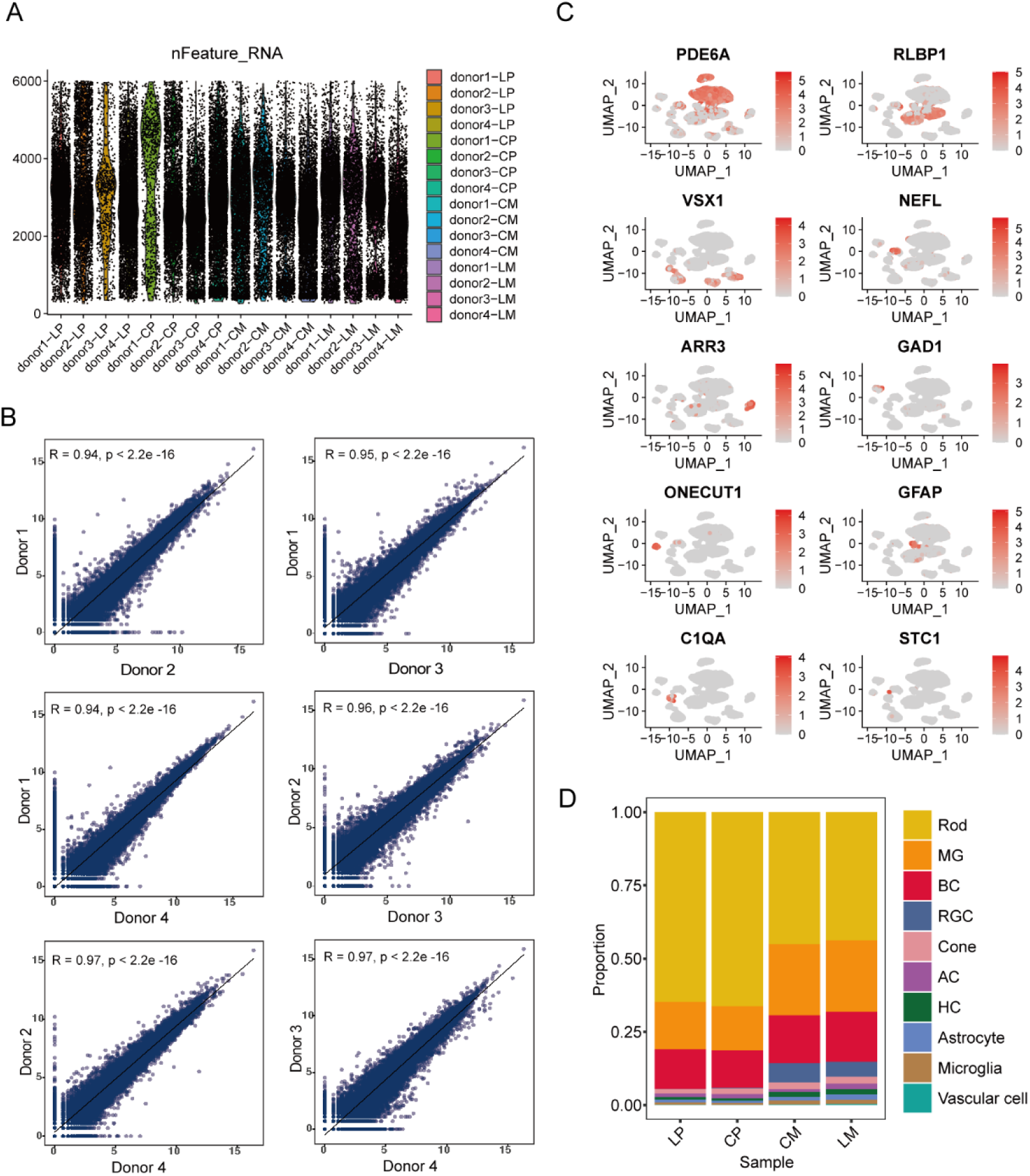
Sequencing depth and biological differences across four donor retinas. **A**. Average number of detected RNA features per cell for each sample. **B**. Correlations of overall gene expression patterns among all donor retinas. **C**. UMAP analysis showing retinal cell markers across different cell types. **D**. Proportions of retinal cell types in different treatment groups.

**Supplementary Figure 3.**
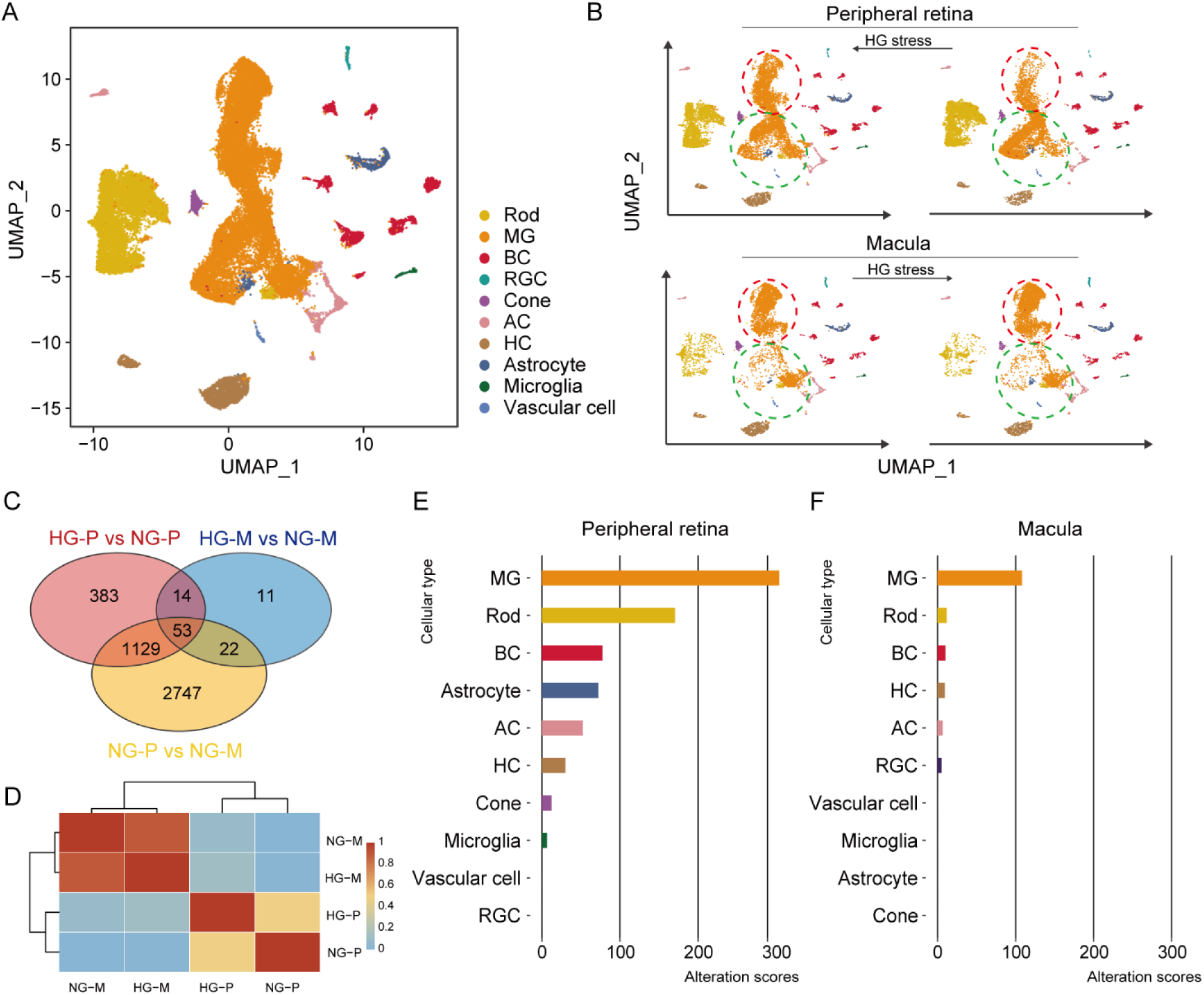
Differences between the macula and peripheral retinas with or without hyperglycaemic stress. To determine if the findings from light-stressed retinas reflect a broader stress response, we conducted a pilot single-cell RNA sequencing (scRNA-seq) study on human maculas (**M**) and peripheral retinas (**P**) cultured *ex vivo* under hyperglycemic (**HG**) stress or normal glucose (**NG**) as a control. We identified 38,761 single cells and 10 retinal cell types from macular and peripheral retinal explants cultured in normal or high glucose conditions. **A**. Uniform manifold approximation and projection (UMAP) visualization of 38,761 single cells from macular and peripheral retinal explants cultured in normal and high glucose conditions. **B**. Two major subtypes of Müller glial (MG) cells (orange) with distinct transcriptomic profiles, one dominant in the human macula (red circle) and the other in the peripheral retina (green circle). **C**. Venn diagram showing the number of significantly differentially expressed genes (DEGs) among treatment groups. **HG-P**: peripheral retina with high glucose; **NG-P**: peripheral retina with normal glucose; **HG-M**: macula with high glucose; **NG-M**: macula with normal glucose. **D**. Heatmap illustrating correlations between the different treatment groups. High correlation (red) in NG-M vs. HG-M represents fewer transcriptional changes; low correlation (yellow) in NG-P vs. HG-P represents significant transcriptomic profile changes. **E** & **F**. Alteration scores of different retinal cell types in the peripheral retina (**E**) and macula (**F**) in response to high glucose. A higher alteration score indicates more significant transcriptional changes.

**Supplementary Figure 4.**
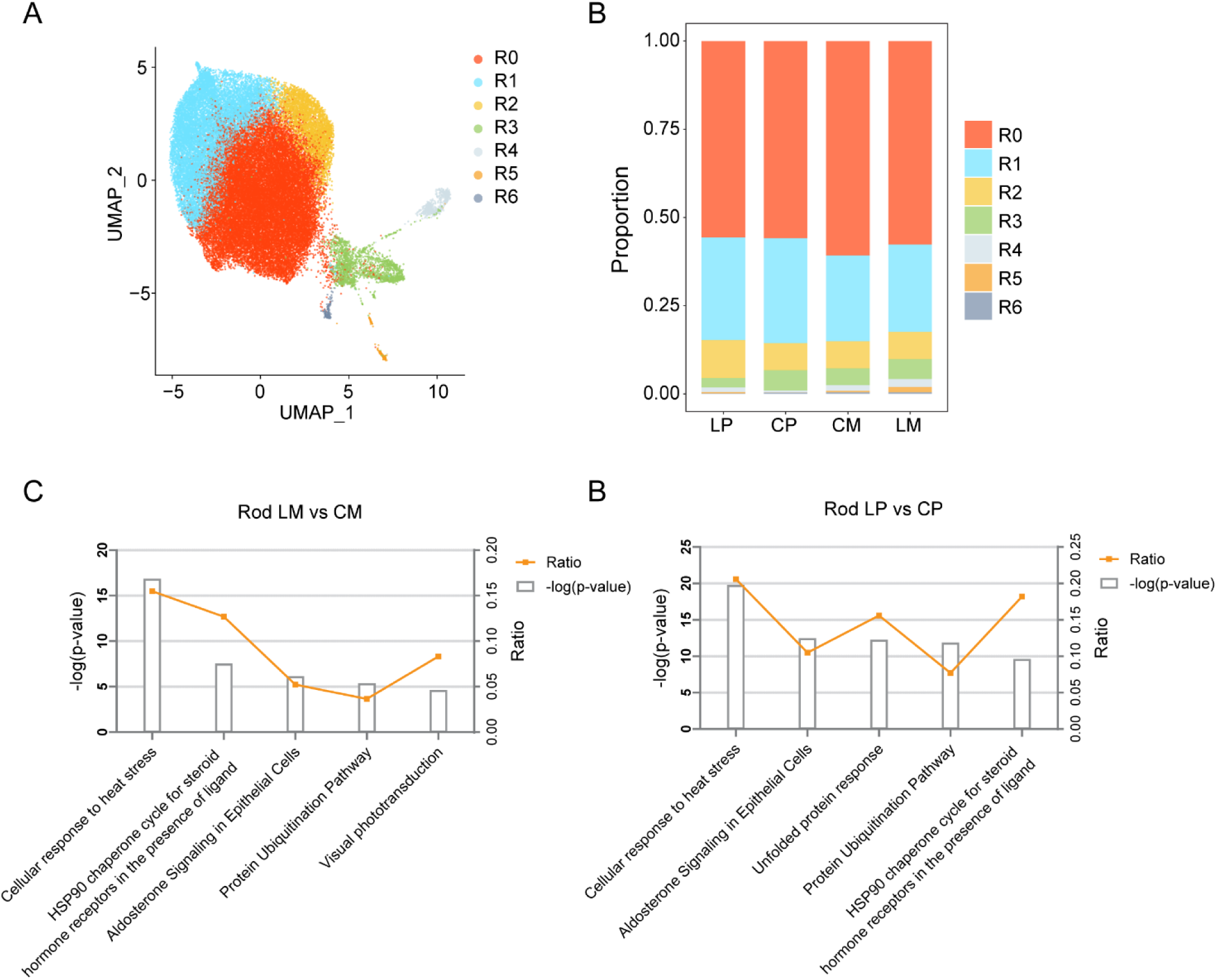
Analysis of rod subtypes and their proportions, along with Ingenuity Pathway Analysis (IPA) comparisons between LM vs. CM and LP vs. CP in rods. **A.** UMAP visualization of rod subtypes. **B**. Proportions of different rod subtypes. **C**. IPA comparing LM and CM within rod subtypes. **D**. IPA comparing LP and CP within rod subtypes.

**Supplementary Figure 5.**
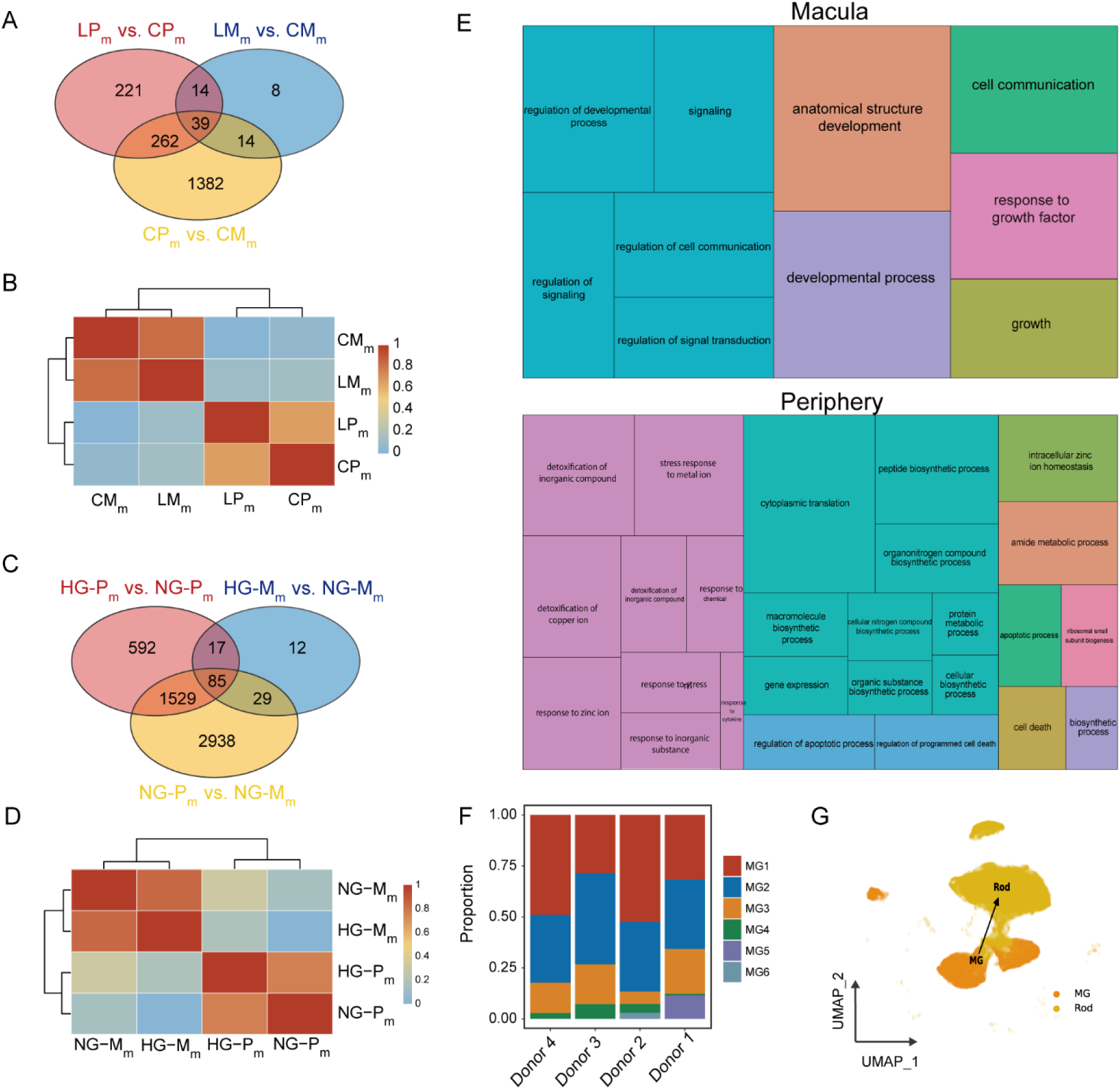
Comparison of transcriptomic changes in Müller cells: Peripheral vs. Macular in response to light and hyperglycaemic stress. We investigated the DEGs between total Müller cells from the macula and peripheral retina after exposure to light stress or hyperglycaemic stress across the four treatment groups. **A**. Venn diagram showing significant DEGs among treatment groups with or without light stress. **B**. Correlation analysis on transcriptomes in all Müller cells of four treatment groups in response to light stress. **LP_m_**: Light-stressed Peripheral Müller cells, **CP_m_**: Control Peripheral Müller cells, **LM_m_**: Light-stressed Macular Müller cells, **CM_m_**: Control Macular Müller cells. **C**. Venn diagram showing significant DEGs among the treatment groups with or without hyperglycaemic stress. **D**. Correlation analysis on transcriptomes in all Müller cells of four treatment groups in response to high glucose. **HG-P_m_**: peripheral Müller cells with high glucose; **NG-P_m_:** peripheral Müller cells with normal glucose; **HG-M_m_**: macular Müller cells with high glucose; **NG- M_m_**: macular Müller cells with normal glucose. **E**. Treemap of GO analysis on biological processes in human macular and peripheral Müller cells. **F**. Proportions of subtypes of Müller cells in different donors. **G**. Partition-based graph abstraction depicting the direction of differentiation from MG to rods.

**Supplementary Figure 6.**
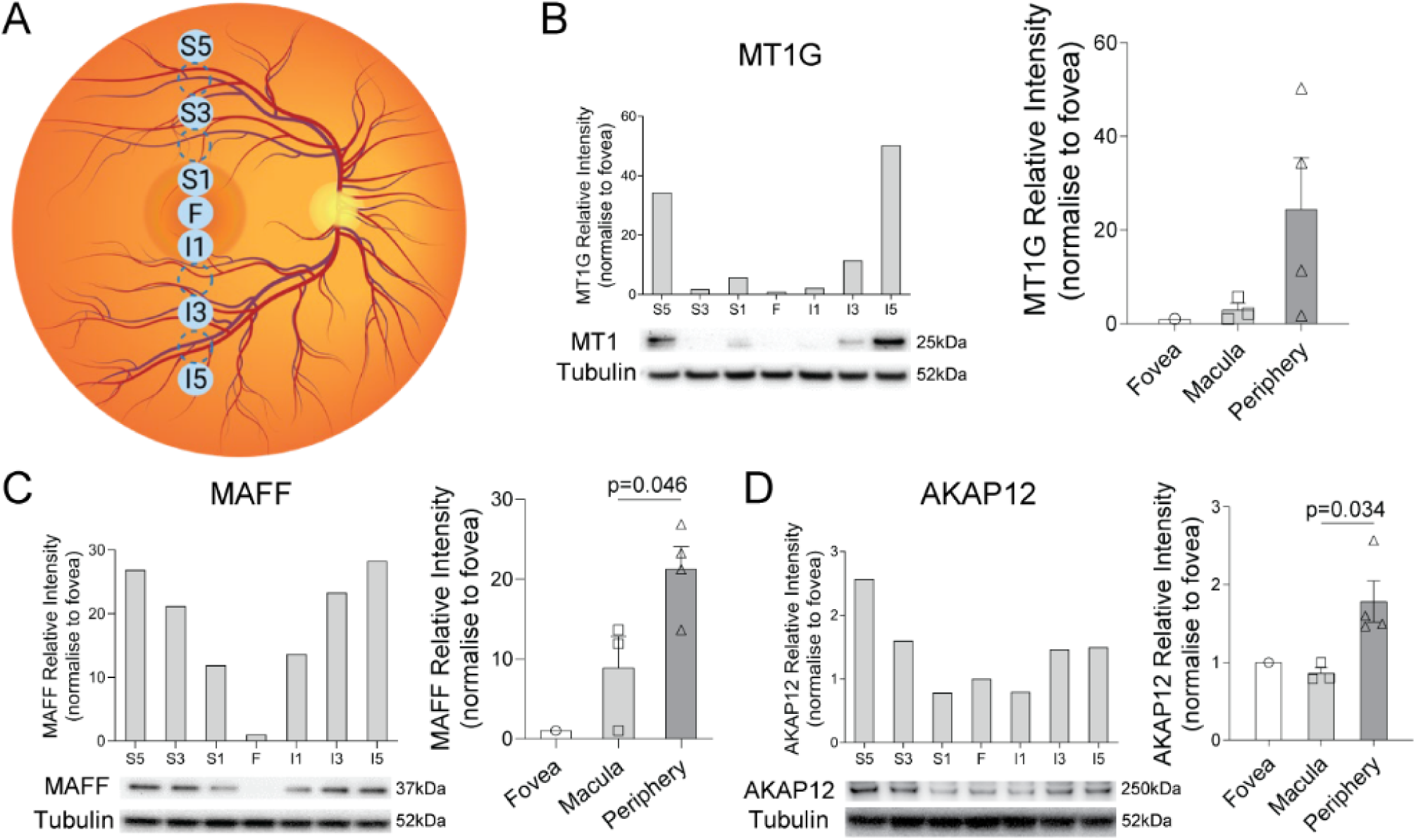
Topographic protein expressions of MT1, MAFF and AKAP12 in human postmortem retina. **A**. Schematic of retinal topographic punches in the macula and peripheral retina. Solid circles ● represent 2mm-diameter retinal punches used for protein validation, while dotted circles ◌ represent 2mm-diameter retinal punches not used for protein validation. **B-D**. Protein expressions of MT1G, MAFF and AKAP12 in different locations of the neural retina. S: superior, I: inferior, F: fovea.

**Supplementary Figure 7.**
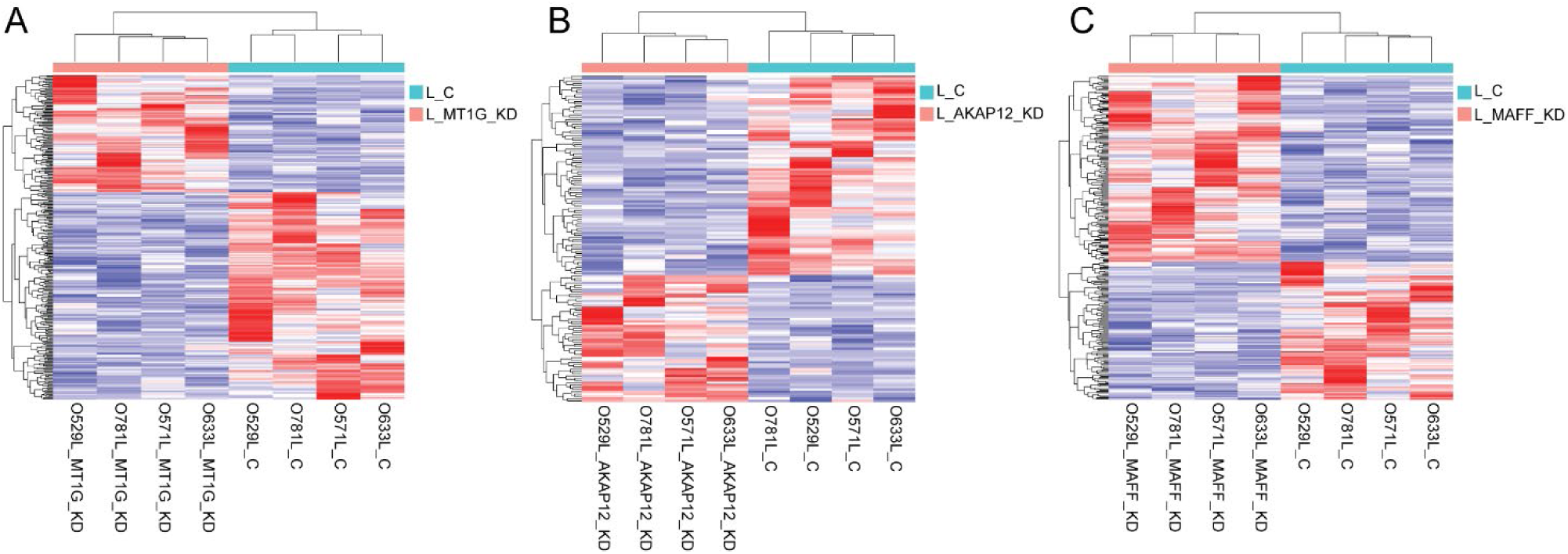
Heatmaps of gene expressions in in human primary Müller cells with AKAP12, MAFF and MT1G siRNA knockdown or the control group under light stress. **A-C.** Heatmaps of gene expression and clustering of individual samples between AKAP12, MAFF and MT1G siRNA knockdown and control groups.

**Supplementary Figure 8.**
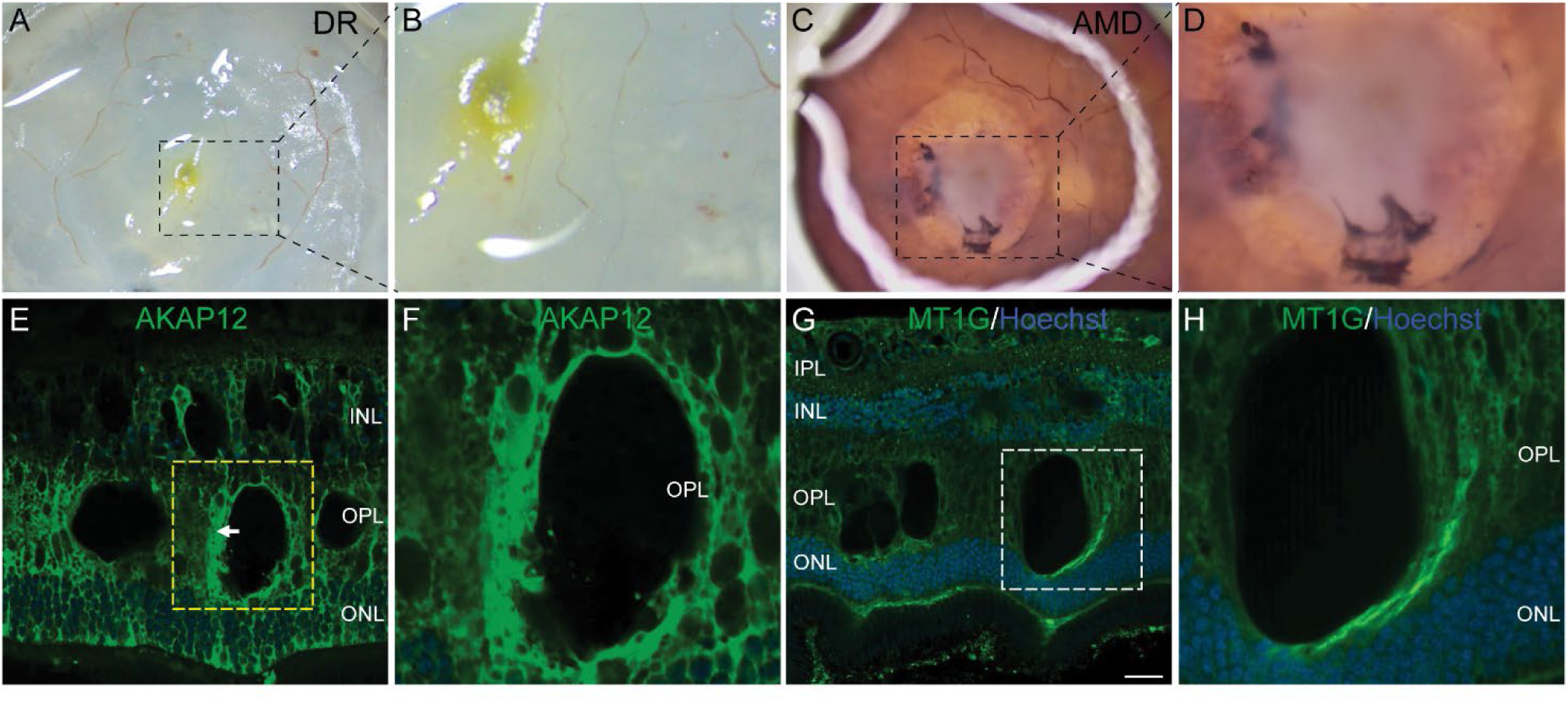
Postmortem retinas with Diabetic Retinopathy (DR) and dry age-related macular degeneration (AMD) and Activation of AKAP12 and MT1 in the retina with DR. **A**. The fundus photo of the retina with DR. **B**. Magnified image in the dotted box of **A**. **C**. The fundus photo of the retina with dry AMD. **D**. Magnified image in the dotted box of **C**. **E**. Immunofluorescent (IF) staining of AKAP12 (green) and Hoechst (blue) on the doner retina with DR. **F**. Magnified image in the yellow dotted box of **E**. **G**. IF staining of MT1 (green) and Hoechst (blue) on the donor retina with DR. **H**. Magnified image in the white dotted boxes of **G.**

**Supplementary Table 1.** Donor Information.

| Donor Number | Sex | Age | Postmortem |
| --- | --- | --- | --- |
| Donor 1 | F | 57y | 18h |
| Donor 2 | F | 63y | 20h |
| Donor 3 | F | 70y | 5h |
| Donor 4 | F | 75y | 20h |

| Selection Criteria |  |
| --- | --- |
| • | post-mortem delay <20h |
| • | less than 75 years old |
| • | no eye conditions, diabetes, jaundice |
| • | no multi-organ failure, head trauma |

## Notes

### Competing Interest Statement

The authors have declared no competing interest.

### Summary of Updates

Data Refinement: We corrected figure numbering and labeling inconsistencies, ensuring better clarity in figures, particularly Figures 2, 3, 5, and 6. New Data on Human Retinal Tissues: To enhance the relevance of our findings, we conducted additional immunofluorescent staining for Muller cell markers (AKAP12, MT1, MAFF) in retinal tissues from donor patients with age-related macular degeneration and diabetic retinopathy. These data further substantiate the translational significance of our work. Expanded Bioinformatics Analysis: We incorporated additional analyses on rod cells from the macula and peripheral retina, highlighting the interplay between Muller cells and rod populations in response to light stress. Clarification of Experimental Details: We provided further explanations regarding the exposure times, intensity of light stress, and the methodologies used in our study to address reviewer concerns.

